# Genomic characterization of endemic and ecdemic non-typhoidal *Salmonella enterica* lineages circulating among animals and animal products in South Africa

**DOI:** 10.1101/2021.05.06.442881

**Authors:** Laura M. Carroll, Rian Pierneef, Masenyabu Mathole, Itumeleng Matle

## Abstract

Non-typhoidal *Salmonella enterica* imposes a significant burden on human and animal health in South Africa. However, very little is known about lineages circulating among animals and animal products in the country on a genomic scale. Here, we used whole-genome sequencing (WGS) to characterize 63 *Salmonella enterica* strains (*n* = 18, 8, 13, and 24 strains assigned to serotypes Dublin, Hadar, Enteritidis, and Typhimurium, respectively) isolated from livestock, companion animals, wildlife, and animal products in South Africa over a 60-year period. Within-serotype phylogenies were constructed using genomes sequenced in this study, as well as publicly available genomes representative of each respective serotype’s (i) global (*n* = 2,802 and 1,569 *S.* Dublin and Hadar genomes, respectively) and (ii) African (*n* = 716 and 343 *S.* Enteritidis and Typhimurium genomes, respectively) population. For *S.* Dublin, the approaches used here identified a largely antimicrobial-susceptible, endemic lineage circulating among humans, animals, and food in South Africa, as well as a lineage that was likely recently introduced from the United States. For *S.* Hadar, multiple South African lineages harboring streptomycin and tetracycline resistance-conferring genes were identified. African *S.* Enteritidis could be primarily partitioned into one largely antimicrobial-susceptible and one largely multidrug-resistant (MDR) clade, with South African isolates confined to the largely antimicrobial-susceptible clade. *S.* Typhimurium strains sequenced here were distributed across the African *S.* Typhimurium phylogeny, representing a diverse range of lineages, including numerous MDR lineages. Overall, this study provides insight into the evolution, population structure, and antimicrobial resistome composition of *Salmonella enterica* in Africa.

**IMPORTANCE:** Globally, *Salmonella enterica* is estimated to be responsible for more than 93 million illnesses and 150,000 deaths annually. In Africa, the burden of salmonellosis is disproportionally high; however, WGS efforts are overwhelmingly concentrated in world regions with lower salmonellosis burdens. While WGS is being increasingly employed in South Africa to characterize *Salmonella enterica*, the bulk of these efforts have centered on characterizing human clinical strains. WGS data derived from non-typhoidal *Salmonella enterica* serotypes isolated from non-human sources in South Africa is extremely limited. To our knowledge, the genomes sequenced here represent the largest collection of non-typhoidal *Salmonella enterica* isolate genomes from non-human sources in South Africa to date. Furthermore, this study provides critical insights into endemic and ecdemic non-typhoidal *Salmonella enterica* lineages circulating among animals, foods, and humans in South Africa and showcases the utility of WGS in characterizing animal-associated strains from a world region with a high salmonellosis burden.

## INTRODUCTION

Livestock, domestic animals, and wildlife can serve as potential reservoirs for non-typhoidal *Salmonella enterica* (1, 2). As a zoonotic foodborne pathogen, *Salmonella enterica* can be transmitted from these animal reservoirs to humans, either via direct contact with infected animals or along the food supply chain (2, 3); however, evolutionary lineages within the *Salmonella enterica* species may vary in terms of their host specificity, geographic distribution, and the severity of illness that they cause in a given host (2, 4). *Salmonella enterica* serotype Typhimurium (*S.* Typhimurium), for example, can infect a broad range of species, while serotype Dublin (*S.* Dublin) is largely adapted to cattle, but can cause rare but frequently invasive infections in humans (5–11).

Due to its importance as a pathogen from both a human and animal health perspective, there is a strong incentive to monitor the evolution and spread of *Salmonella enterica* in animals and animal products (12, 13). Furthermore, there has been growing concern that *Salmonella enterica* can acquire antimicrobial resistance (AMR) determinants in livestock environments, which can make infections in humans and animals more difficult and costly to treat (14, 15). To this end, whole-genome sequencing (WGS) is being increasingly employed to characterize *Salmonella enterica* from animals (e.g., livestock, companion animals, and wildlife) and animal products, as WGS can not only replicate many important microbiological assays *in silico* (e.g., prediction of serotype, AMR), but provide additional data that can be used to characterize isolates (e.g., identification of genome-wide single nucleotide polymorphisms [SNPs], core- and whole-genome multi-locus sequence typing [MLST], pan-genome characterization) (16–19).

In South Africa, the bulk of *Salmonella enterica* WGS efforts have focused on characterizing human clinical strains associated with illnesses and/or outbreaks (20–25). WGS-based studies querying *Salmonella enterica* strains isolated from non-human sources in South Africa are limited (26), and little is known regarding which lineages are circulating among animals in the country (27). Here, we used WGS to characterize 63 South African *Salmonella enterica* strains isolated from animals and animal products over the course of 60 years (i.e., between 1960 and 2019). Using phylogenomic approaches, we characterized the isolates sequenced here within the context of publicly available genomes representative of the global (for *S.* Dublin and *S.* Hadar) and African (for *S.* Enteritidis and *S.* Typhimurium) *Salmonella enterica* populations. The results presented here will provide critical insights into the evolution, population structure, and AMR dynamics of *Salmonella enterica* in Africa.

## RESULTS

### Four serotypes are represented among the animal-associated South African *Salmonella enterica* strains sequenced here

A total of 63 *Salmonella enterica* strains were isolated from animals and animal products in South Africa and underwent WGS (Supplemental Table S1). All isolates underwent *in silico* serotyping using both (i) SISTR (using its core-genome MLST [cgMLST] approach) and (ii) SeqSero2 (Supplemental Table S1); serotypes assigned using both methods were identical for all isolates (63 of 63 isolates, 100%; Supplemental Table S1). Furthermore, genomes of all isolates sequenced here clustered among publicly available *Salmonella* genomes assigned to their respective serotypes (28), with no observed polyphyly within serotypes among isolates sequenced here (Figure 1; note that all *S.* Hadar genomes sequenced here clustered among a genome previously serotyped as *S.* Istanbul, which was serotyped as *S.* Hadar *in silico* using both SISTR and SeqSero2).

**Figure 1.**
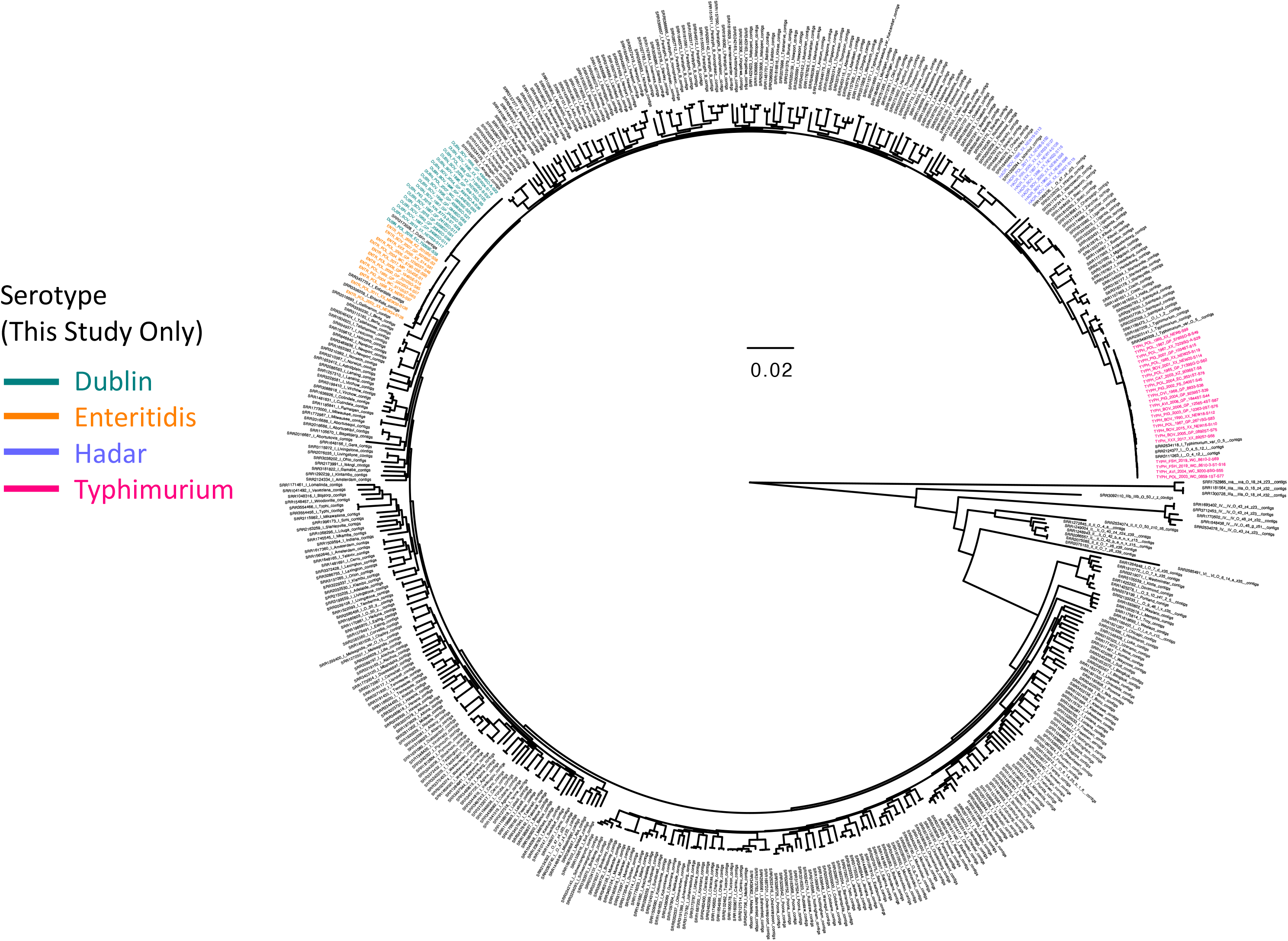
Maximum likelihood phylogeny constructed using core SNPs identified among 505 *Salmonella* isolate genomes. Publicly available genomes are denoted by black tip labels (*n* = 442), while genomes of strains isolated in conjunction with this study are denoted by tip labels colored by serotype (*n* = 63). The phylogeny is rooted at the midpoint with branch lengths reported in substitutions per site. Core SNPs were identified among all genomes using kSNP3, while the phylogeny was constructed and annotated using IQ-TREE and FigTree v. 1.4.4, respectively.

Four serotypes were represented among isolates sequenced in this study: *S.* Dublin, *S.* Hadar, *S.* Enteritidis, and *S.* Typhimurium, assigned to 18, 8, 13, and 24 isolates, respectively (Figure 1 and Supplemental Table S1). Strains were isolated from bovine sources (from feces, meat, or organs; *n* = 25), poultry (from feces, meat, or organs; *n* = 22), swine (from feces, meat, or organs; *n* = 6), unknown sources (*n* = 3), fish (from food products; *n* = 2), avian sources (feces from each of an ostrich and a pigeon; *n* = 2), a rhinoceros (*n* = 1), ovine sources (from feces; *n* = 1), and from a cat (from feces; *n* = 1; Supplemental Table S1). Strains were isolated from one of six provinces in South Africa: Gauteng (*n* = 27), Western Cape (*n* = 7), KwaZulu-Natal (*n* = 2), Eastern Cape (*n* = 2), North-West (*n* = 1), Mpumalanga (*n* = 1), and Free State (*n* = 1); the provinces from which an additional 22 strains were isolated were unknown (Supplemental Table S1).

### AMR in *Salmonella enterica* isolated from animals and animal products in South Africa is acquired sporadically

The 63 *Salmonella* genomes sequenced here underwent *in silico* AMR/stress response determinant, plasmid replicon, and virulence factor detection (Figure 2 and Supplemental Figures S1 and S2). In total, 59 different AMR/stress response determinants were detected among the 63 isolates, with 18 unique AMR/stress response determinant presence/absence profiles observed (based on AMR/stress response determinants detected using AMRFinderPlus; Figure 2) (29). The number of different AMR/stress response determinants detected per genome ranged from five to 24; nearly two-thirds of all genomes sequenced in the current study (40 of 63, 63.5%) harbored six AMR/stress response determinants (the median per genome) or less (Figure 2). Six “core” AMR/stress response determinants (*asr, golS, golT, mdsA, mdsB, sinH*) were observed in over 90% of the isolates sequenced here (59 of 63 isolates; 93.7%), four of which were detected in all 63 isolates (*asr, golS, golT, sinH*; Figure 2). The remaining 53 AMR/stress response determinants were detected in less than 20% of the genomes sequenced here; 46 of these (46 of 59 total unique AMR determinants, 78.0%) were present only sporadically and were detected in two or fewer genomes (Figure 2).

**Figure 2.**
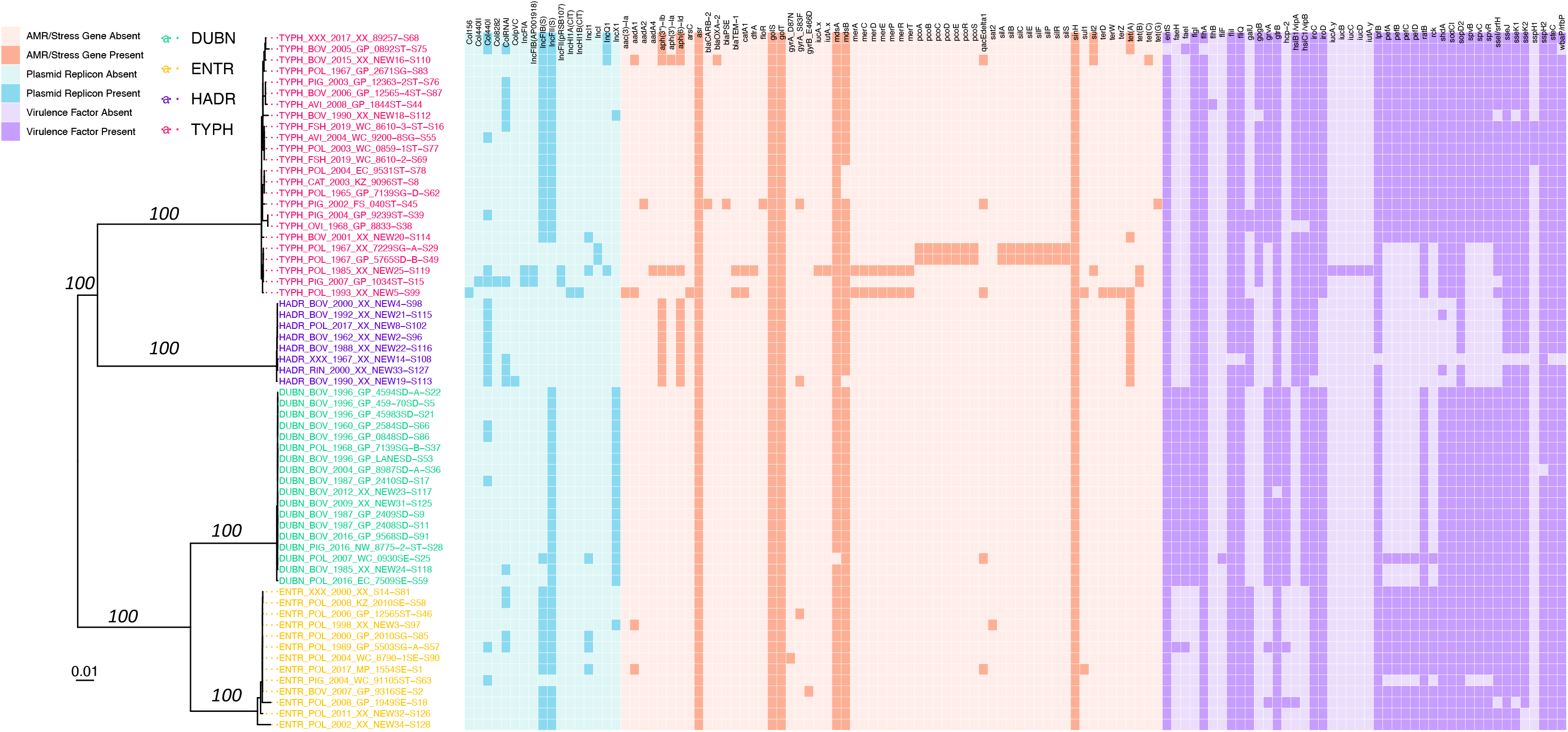
Maximum likelihood phylogeny constructed using core SNPs identified among the genomes of 63 *Salmonella* strains isolated in conjunction with this study. Tip label colors denote isolate serotypes, and branch labels denote ultrafast bootstrap support percentages out of 1,000 replicates (selected for readability). The heatmap to the right of the phylogeny denotes the presence and absence of (i) plasmid replicons (blue), (ii) antimicrobial resistance (AMR) and stress response determinants (orange), and (iii) variably detected virulence factors (purple) in each genome. The phylogeny is rooted at the midpoint with branch lengths reported in substitutions per site. Core SNPs were identified among all genomes using kSNP3. Plasmid replicons were identified using ABRicate and the PlasmidFinder database, using minimum identity and coverage thresholds of 80 and 60%, respectively. AMR and stress response determinants were identified using AMRFinderPlus. Virulence factors were identified using ABRicate and VFDB, using minimum identity and coverage thresholds of 70 and 50%, respectively. Virulence factors detected in all genomes were excluded for readability (Supplemental Table S2). The phylogeny was constructed and annotated using IQ-TREE and bactaxR/ggtree, respectively. DUBN, *S.* Dublin; ENTR, *S.* Enteritidis; HADR, *S.* Hadar; TYPH, *S.* Typhimurium.

In total, 17 different plasmid replicons were identified among all 63 genomes, representing 22 unique plasmid replicon presence/absence profiles (detected using ABRicate, the PlasmidFinder database, and minimum nucleotide identity and coverage thresholds of 80 and 60%, respectively; Figure 2) (30). Genomes harbored one to seven different plasmid replicons, with a median of two per genome (Figure 2). Two plasmid replicons, IncFIB(S) and IncFII(S), were detected in over half of all genomes sequenced here (detected in 32 and 49 of 63 genomes, 50.8% and 77.8%, respectively; Figure 2). Over half of all plasmid replicons (10 of 17 unique plasmid replicons; 58.8%) were detected in two or fewer genomes (Figure 2).

Additionally, a total of 181 different virulence factors were identified among the 63 genomes, with 24 unique virulence factor presence/absence profiles represented (detected using ABRicate, the Virulence Factor Database [VFDB], and minimum nucleotide identity and coverage thresholds of 70 and 50%, respectively; Figure 2 and Supplemental Table S2). Genomes harbored 146 to 171 different virulence factors, with a median of 165 (Figure 2 and Supplemental Table S2). Over 75% of all unique virulence factors detected among the isolates sequenced in this study were present in all genomes (137 of 181 unique virulence factors, 75.7%; Supplemental Table S2). Only 13 virulence factors were detected in fewer than half of the genomes sequenced here (Figure 2).

### A largely antimicrobial-susceptible *S.* Dublin ST10 lineage circulating in South Africa encompasses isolates from livestock, food, and human sources

A maximum likelihood (ML) phylogeny constructed using the 18 South African sequence type 10 (ST10) *S.* Dublin isolates sequenced here, plus 2,784 publicly available ST10 *S.* Dublin genomes, partitioned the vast majority of genomes (2,738 of 2,802 genomes, 97.7%) into two major *S.* Dublin ST10 clades (Figure 3 and Supplemental Figure S3), which is consistent with previous observations (31). Referred to hereafter as “*S.* Dublin Major Clade I” and “*S.* Dublin Major Clade II”, the two major clades encompassed 1,787 and 951 genomes, respectively (Figure 3 and Supplemental Figure S3). While both major clades encompassed strains isolated from Asia, Europe, North America, and South America, the vast majority of North American ST10 *S.* Dublin belonged to Major Clade I (1,641 of 1,656 *S.* Dublin ST10 strains from North America, 99.1%; Figure 3 and Supplemental Figure S3). Members of Major Clade I shared a most recent common ancestor (MRCA) dated to circa 1959 (95% confidence interval [CI] of [1452.88, 1959.00]; Figure 3 and Supplemental Figure S3). Notably, multi-drug resistant (MDR) *S.* Dublin, which often possess IncA/C2 plasmids and acquired AMR determinants that confer resistance to aminoglycosides, beta-lactams, phenicols, sulfonamides, and tetracyclines (31), were almost exclusively confined to a large, primarily North American subclade within *S.* Dublin Major Clade I (referred to hereafter as the “*S.* Dublin Large Subclade”; Figure 3 and Supplemental Figure S3). Conversely, members of *S.* Dublin Major Clade II shared a MRCA dated to circa 1945 (95% CI [1274.31, 1985.00]), primarily contained European isolates (893 of 951 Major Clade II genomes, 93.9%), and largely did not possess any acquired AMR determinants (Figure 3 and Supplemental Figure S3).

**Figure 3.**
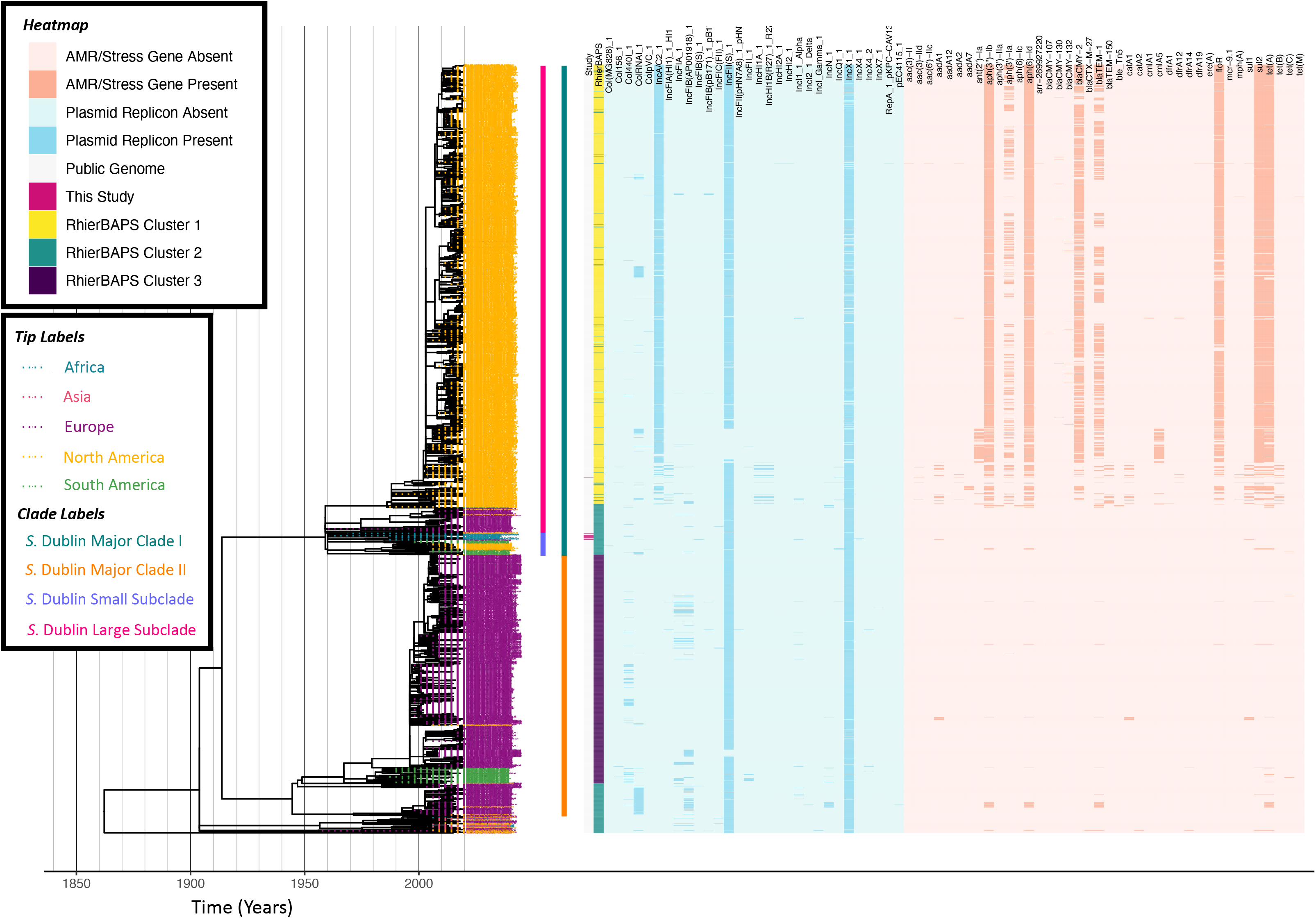
Maximum likelihood phylogeny constructed using core SNPs identified among 2,802 *S.* Dublin genomes (2,784 publicly available genomes, plus 18 sequenced here). Tip label colors denote the continent from which each strain was reported to have been isolated. Clade labels denote major clades assigned in this study and are shown to the right of tip labels. The heatmap to the right of the phylogeny denotes: (i) whether an isolate was sequenced in conjunction with this study (dark pink) or not (gray; “Study”); (ii) level 1 cluster assignments obtained using RhierBAPS (“RhierBAPS”); the presence and absence of (iii) plasmid replicons (blue) and (iv) antimicrobial resistance (AMR) determinants (orange). The phylogeny was rooted and time-scaled using LSD2, with branch lengths reported in years (X-axis). Core SNPs were identified among all genomes using Parsnp. AMR determinants were identified using ABRicate, the NCBI AMR determinant database, and minimum identity and coverage thresholds of 75 and 50%, respectively. Plasmid replicons were identified using ABRicate and the PlasmidFinder database, using minimum identity and coverage thresholds of 80 and 60%, respectively. The phylogeny was constructed and annotated using IQ-TREE and bactaxR/ggtree, respectively.

All 18 South African *S.* Dublin isolates sequenced in this study belonged to *S.* Dublin Major Clade I (Figure 3 and Supplemental Figure S3); however, 17 of the 18 isolates clustered together within a small subclade of Major Clade I (referred to hereafter as the “*S.* Dublin Small Subclade”; Figure 4), while the remaining isolate clustered among isolates in the *S.* Dublin Large Subclade (Supplemental Figure S4). Notably, within the *S.* Dublin Small Subclade, the 17 animal- and animal product-associated South African isolates sequenced here clustered among all seven publicly available *S.* Dublin genomes from South Africa, all of which were reported to have been isolated from human sources (Figure 4). This well-supported South African-specific *S.* Dublin lineage (referred to hereafter as the “South African *S.* Dublin Clade”), which contained animal-, animal product-, and human-associated strains isolated over a span of 60 years (i.e., 1960-2020) from the Gauteng, Eastern Cape, Western Cape, and North-West provinces, was predicted to share a common ancestor dated to circa 1960 (95% CI [1496.07, 1960.00], 98% UltraFast Bootstrap Support; Figure 4). Members of this South African lineage, like the *S.* Dublin Small Subclade more broadly, were largely pan-susceptible, with AMR determinants detected only sporadically; a single strain, isolated in 2007 from poultry meat in the Western Cape province (FOO_2007_SouthAfrica_WesternCape_AF_0930SE−S25), possessed streptomycin resistance gene *aadA1* and sulfonamide resistance gene *sul1* (Figure 4). Taken together, these results indicate that a largely AMR-susceptible South African-specific *S.* Dublin lineage has been circulating among animals, foods, and humans in the country for decades.

**Figure 4.**
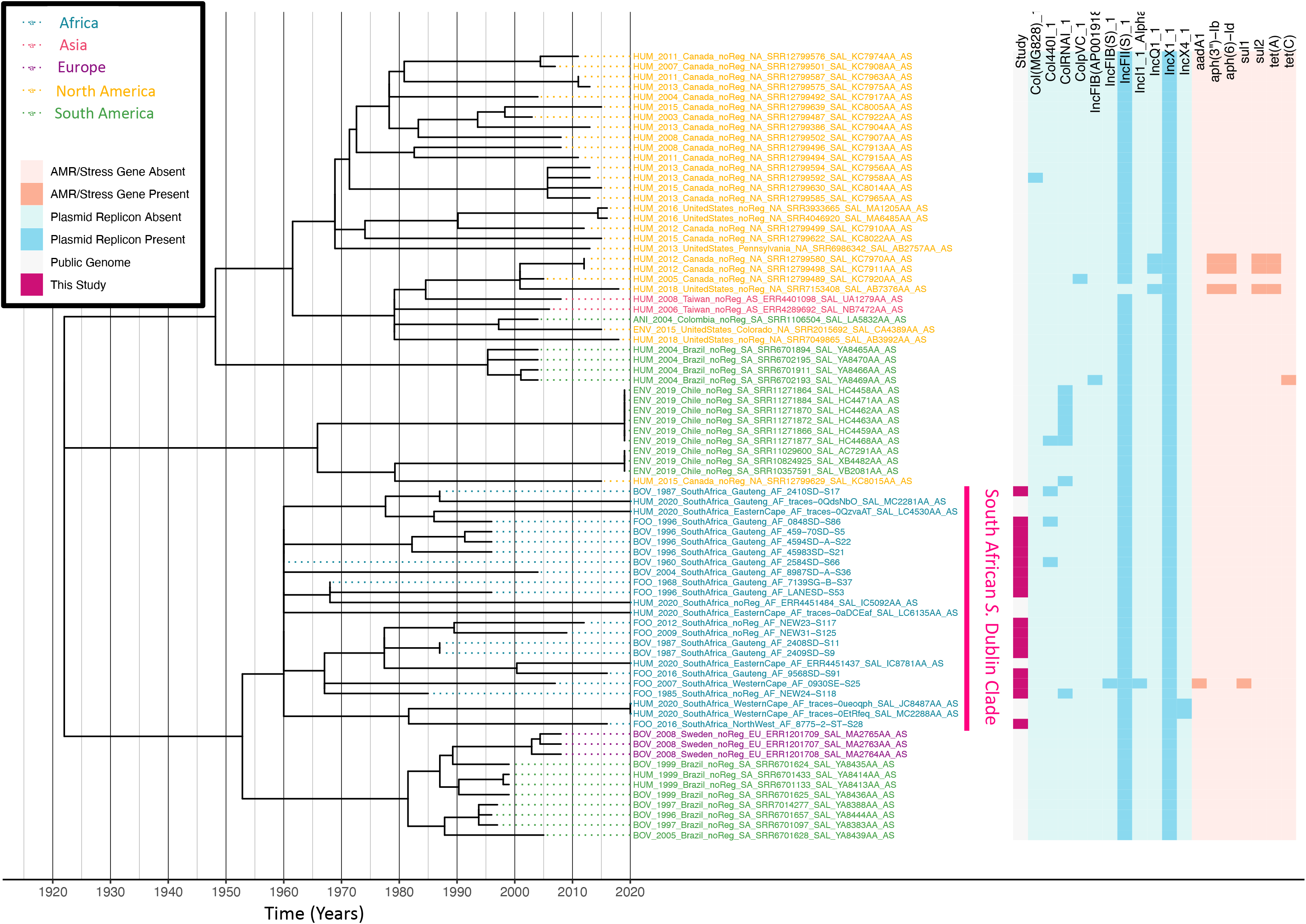
Maximum likelihood phylogeny constructed using core SNPs identified among 78 *S.* Dublin genomes within the *S.* Dublin Small Subclade (61 publicly available genomes, plus 17 sequenced here). Tip label colors denote the continent from which each strain was reported to have been isolated. A pink clade label to the right of the tip labels denotes a clade of South African isolates, which encompasses 17 of the 18 *S.* Dublin isolates sequenced in this study, plus seven publicly available South African isolates. The heatmap to the right of the phylogeny denotes: (i) whether an isolate was sequenced in conjunction with this study (dark pink) or not (gray; “Study”); the presence and absence of (ii) plasmid replicons (blue) and (iii) antimicrobial resistance (AMR) determinants (orange). The phylogeny was rooted and time-scaled using LSD2, with branch lengths reported in years (X-axis). Core SNPs were identified among all genomes using Parsnp. AMR determinants were identified using ABRicate, the NCBI AMR determinant database, and minimum identity and coverage thresholds of 75 and 50%, respectively. Plasmid replicons were identified using ABRicate and the PlasmidFinder database, using minimum identity and coverage thresholds of 80 and 60%, respectively. The phylogeny was constructed and annotated using IQ-TREE and bactaxR/ggtree, respectively.

Only one South African *S.* Dublin genome was not a member of the South African *S.* Dublin Clade within the *S.* Dublin Small Subclade (Supplemental Figure S4). This strain (i.e., FOO_2016_SouthAfrica_EasternCape_AF_7509SE-S59), which was isolated in 2016 from poultry meat in South Africa’s Eastern Cape province, clustered among North American isolates in the *S.* Dublin Large Subclade (Supplemental Figure S4). This strain most closely resembled a bovine-associated strain from California isolated in 2004, and the two shared a common ancestor circa 2004 (95% CI [1973.4, 2004.00]; Supplemental Figure S4). Notably, despite clustering among MDR *S.* Dublin strains from North America, neither of these strains harbored any acquired AMR genes, nor did they harbor the IncA/C2 plasmid characteristic of MDR *S.* Dublin from the United States (Supplemental Figure S4). These results indicate that a separate *S.* Dublin lineage may have only recently been introduced into South Africa from North America, a hypothesis that was further supported by subsequent investigation into the origin of the isolate: the poultry meat from which strain FOO_2016_SouthAfrica_EasternCape_AF_7509SE-S59 was isolated had been imported from North America and sold in a supermarket in South Africa’s Eastern Cape province.

All 18 *S.* Dublin isolates sequenced in this study, as well as all seven publicly available South African *S.* Dublin genomes, were members of *S.* Dublin Major Clade I (Figure 3 and Supplemental Figure S3). These 25 South African genomes, 24 of which formed a well-supported subclade within Major Clade I, were the only African genomes detected in *S.* Dublin Major Clade I (Figure 3 and Supplemental Figures S3-S4). *S.* Dublin Major Clade II did not contain any African genomes (Figure 3). However, 18 genomes from the African continent were among the few genomes (i.e., 64 of 2,802 *S.* Dublin genomes, 2.3%) that fell outside of the two major *S.* Dublin clades (Figure 3). These genomes were reported to have been derived from strains isolated from animals, food, and humans in Ethiopia, Gambia, Nigeria, and Benin, and none harbored any acquired AMR genes (Figure 3); interestingly, they clustered among human-associated genomes from Asia (i.e., Taiwan), Europe (i.e., France and the United Kingdom), and North America (i.e., Canada and the United States), forming a 52-genome, well-supported clade with a common ancestor dated to circa 1957 (95% CI [1142.96, 2003.00], 100% UltraFast Bootstrap Support; Figure 3).

### South Africa harbors multiple *S.* Hadar ST33 lineages with streptomycin and tetracycline resistance-conferring genes

A ML phylogeny was constructed using the eight South African ST33 *S.* Hadar isolates sequenced here, plus 1,561 publicly available ST33 *S.* Hadar genomes (Figure 5). Notably, the majority of *S.* Hadar genomes harbored AMR genes *aph(3’’)-Ib* and *aph*(*6*)*-Id* (*n* = 1,314 and 1,347 of 1,569 *S.* Hadar genomes, 83.7% and 85.9%, respectively; Figure 5). Also known as *strA* and *strB*, respectively, *aph(3’’)-Ib* and *aph*(*6*)*-Id* confer resistance to streptomycin. The majority of *S.* Hadar genomes additionally harbored *tet(A)*, which confers resistance to tetracycline (*n* = 1,320 of 1,569 *S.* Hadar genomes, 84.1%; Figure 5). All eight *S.* Hadar strains sequenced in this study, which were derived from strains isolated between 1962 and 2017, were among the strains that harbored all of streptomycin resistance-conferring *aph(3’’)-Ib* and *aph(6)-Id* and tetracycline resistance-conferring *tet(A)* (Figure 5).

**Figure 5.**
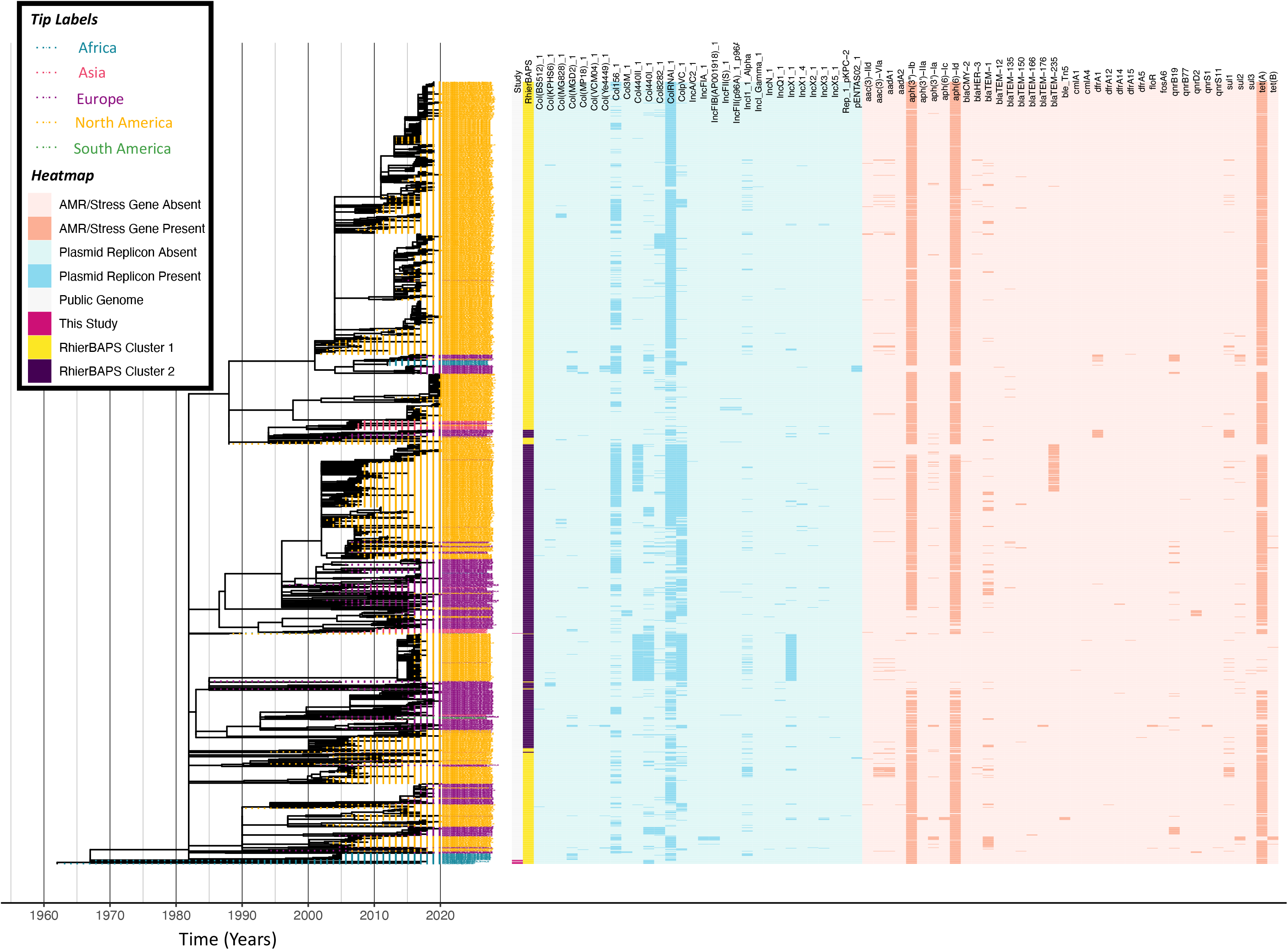
Maximum likelihood phylogeny constructed using core SNPs identified among 1,569 *S.* Hadar genomes (1,561 publicly available genomes, plus eight sequenced here). Tip label colors denote the continent from which each strain was reported to have been isolated. The heatmap to the right of the phylogeny denotes: (i) whether an isolate was sequenced in conjunction with this study (dark pink) or not (gray; “Study”); (ii) level 1 cluster assignments obtained using RhierBAPS (“RhierBAPS”); the presence and absence of (iii) plasmid replicons (blue) and (iv) antimicrobial resistance (AMR) determinants (orange). The phylogeny was rooted and time scaled using LSD2, with branch lengths reported in years (X-axis). Core SNPs were identified among all genomes using Parsnp. AMR determinants were identified using ABRicate, the NCBI AMR determinant database, and minimum identity and coverage thresholds of 75 and 50%, respectively. Plasmid replicons were identified using ABRicate and the PlasmidFinder database, using minimum identity and coverage thresholds of 80 and 60%, respectively. The phylogeny was constructed and annotated using IQ-TREE and bactaxR/ggtree, respectively.

Seven of eight *S.* Hadar genomes sequenced in this study clustered at or near the tree root, which was dated to circa 1962 (95% CI [1571.93, 1962.00]). These seven South African strains, which had been isolated between 1962 and 2017 from bovine sources (feces and meat), poultry (meat), a rhinoceros, and an unknown source, were most closely related to a publicly available genome of a *S.* Hadar strain isolated in 2018 from chicken in South Africa (Figure 5).

The remaining isolate sequenced in this study (i.e., BOV_1990_XX_ARCZA_NEW19-S113) was relatively distantly related to the other South African isolates sequenced here (Figure 5). Isolated from bovine feces in 1990, this strain was most closely related to a *S.* Hadar strain isolated from the spleen of a dog (*Canis lupus familiaris*) in the United States in 1988; however, these strains were relatively distant, sharing a common ancestor that existed circa 1982 (95% CI [1705.27, 1985.00]; Figure 5). While it is unclear exactly when this particular lineage was introduced into South Africa, these results indicate that multiple *S.* Hadar lineages have circulated in livestock populations in the country.

### One largely antimicrobial-susceptible clade and one largely MDR clade are represented among *S.* Enteritidis ST11 from Africa

A ML phylogeny was constructed using the 13 South African ST11 (*n* = 12) and ST366 (*n* = 1) *S.* Enteritidis isolates sequenced here, plus (i) 697 publicly available ST11 *S.* Enteritidis genomes of strains isolated from the African continent and (ii) all publicly available ST366 *S.* Enteritidis genomes (*n* = 10; Figure 6 and Supplemental Figure S5).

**Figure 6.**
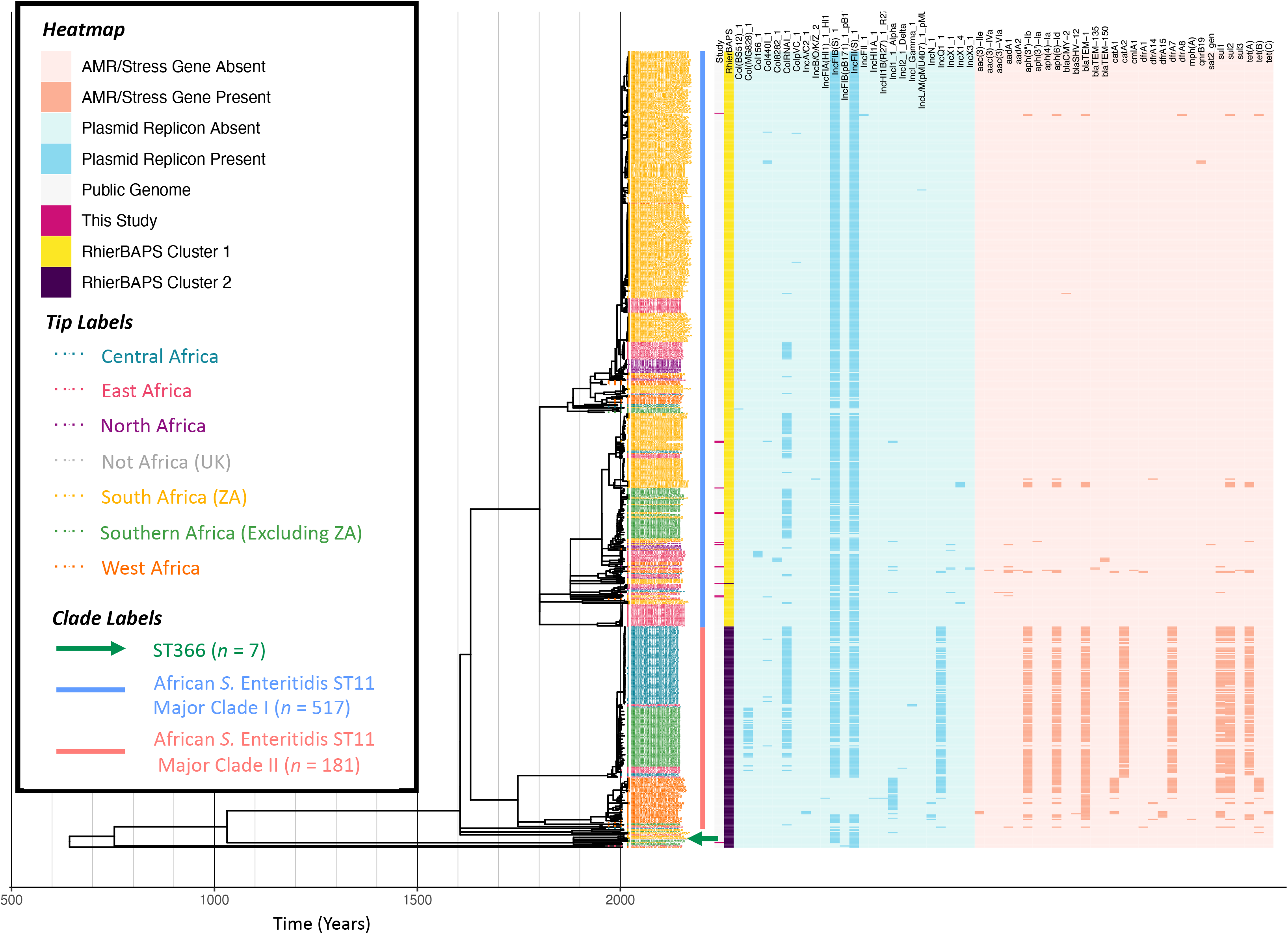
Maximum likelihood phylogeny constructed using core SNPs identified among 716 African *S.* Enteritidis genomes (703 publicly available genomes, plus 13 sequenced here). Tip label colors denote the region/country from which each strain was reported to have been isolated (based on African regions as defined by the African Union, 25 April 2021). Clade labels shown to the right of the phylogeny tip labels denote major clades discussed in the main text. The heatmap to the right of the phylogeny denotes: (i) whether an isolate was sequenced in conjunction with this study (dark pink) or not (gray; “Study”); (ii) level 1 cluster assignments obtained using RhierBAPS (“RhierBAPS”); the presence and absence of (iii) plasmid replicons (blue) and (iv) antimicrobial resistance (AMR) determinants (orange). The phylogeny was rooted and time-scaled using LSD2, with branch lengths reported in years (X-axis). Core SNPs were identified among all genomes using Parsnp. AMR determinants were identified using ABRicate, the NCBI AMR determinant database, and minimum identity and coverage thresholds of 75 and 50%, respectively. Plasmid replicons were identified using ABRicate and the PlasmidFinder database, using minimum identity and coverage thresholds of 80 and 60%, respectively. The phylogeny was constructed and annotated using IQ-TREE and bactaxR/ggtree, respectively.

Notably, one strain sequenced here (i.e., POL_2002_XX_NEW34-S128), isolated from poultry meat in 2002, was assigned to ST366. Currently, there are only 15 ST366 genomes that are publicly available for download, six of which have a known collection year and isolation source and meet the quality standards used in this study (via Enterobase, accessed 18 February 2021). This can be contrasted with ST11, of which there are 50,755 publicly available genomes (via Enterobase, accessed 18 February 2021). The ST366 strain sequenced here was a member of a well-supported clade (100% UltraFast Bootstrap support), which contained eight additional publicly available genomes that shared a common ancestor dated to circa 1885 (95% CI [809.09, 2002.00]; Figure 6 and Supplemental Figure S5). In addition to the poultry-associated ST366 strain sequenced here, this clade contained all six publicly available ST366 genomes, which were all isolated from human sources in South Africa (*n* = 3), Zambia (*n* = 2), and the United Kingdom (*n* = 1); additionally, this clade contained two ST11 genomes of strains isolated from humans in Malawi (Figure 6 and Supplemental Figure S5). None of the genomes in this clade harbored any known AMR determinants (Figure 6 and Supplemental Figure S5). Interestingly, the ST366 isolate from the United Kingdom is the only publicly available ST366 strain from outside of Africa (via Enterobase, accessed 18 February 2021), indicating that this particular ST may have a geographic association.

The remaining 12 *S.* Enteritidis strains sequenced in this study were assigned to ST11 and were confined to a large, well-supported (100% UltraFast Bootstrap support) 517-isolate clade (referred to hereafter as “African *S.* Enteritidis ST11 Major Clade I”), which shared a common ancestor dated to circa 1551-1801 (depending on the tree root/isolate set used in Figure 6 and Supplemental Figure S5, respectively; 95% CI [-639.57, 1955.0] and [738.65, 1955.00], respectively). Notably, isolates within African *S.* Enteritidis ST11 Major Clade I were largely pan-susceptible and acquired AMR determinants only sporadically; among the 12 ST11 isolates sequenced here, only three possessed AMR genes (Figure 6 and Supplemental Figure S5).

Overall, we found that the South African ST11 genomes sequenced in this study belonged to a largely antimicrobial-susceptible lineage, which showcased AMR only sporadically. This can be contrasted with a second major clade comprising 181 *S.* Enteritidis genomes (i.e., African *S.* Enteritidis ST11 Major Clade II; Figure 6); the majority of isolates in this clade were predicted to be MDR, as they possessed AMR genes conferring resistance to beta-lactams (*bla_TEM-1_*), streptomycin (*aph(3’’)-Ib, aph(6)-Id*), sulfonamides (*sul1, sul2*), chloramphenicol (*catA2*), trimethoprim (*dfrA7*), and tetracycline (*tet(A)*; Figure 6). Unlike African *S.* Enteritidis ST11 Major Clade I, which encompassed 340 South African isolates, no Major Clade II isolates were found in South Africa (Figure 6); rather, African *S.* Enteritidis ST11 Major Clade II primarily included isolates from the Democratic Republic of the Congo (DRC; *n* = 72) and Malawi (*n* = 55), as well as from Senegal and Mali (*n* = 12 each), Nigeria (*n* = 10), Kenya (*n* = 7), Burkina Faso (*n* = 5), Rwanda and Guinea (*n* = 2), the Central African Republic (CAR), Congo, Ivory Coast, and Madagascar (*n* = 1 each; Figure 6).

### South Africa harbors numerous antimicrobial susceptible and MDR *S.* Typhimurium lineages

A ML phylogeny was constructed using the 24 South African ST19 (*n* = 23) and ST34 (*n* = 1) *S.* Typhimurium isolates sequenced here, plus publicly available *S.* Typhimurium genomes of strains isolated from the African continent assigned to (i) ST19 (*n* = 315) and (ii) ST34 (*n* = 4; Figure 7 and Supplemental Text). The 24 *S.* Typhimurium strains sequenced in this study were distributed across the African *S.* Typhimurium phylogeny, representing a diverse range of lineages, and eight (33.3%) possessed one or more AMR genes (Figure 7 and Supplemental Text). Notably, some African *S.* Typhimurium lineages were distributed across the African continent, while others were strongly associated with a particular region/country (Figure 7 and Supplemental Text). When compared to genomes from a previous study of *S.* Typhimurium from New York State that have been shown to be representative of the human- and bovine-associated *S.* Typhimurium population in the United States as a whole (32), only five of 24 *S.* Typhimurium strains sequenced here (20.8%) shared a common ancestor with one or more New York State strains after 1900 (Supplemental Figure S6 and Supplemental Text). This indicates that many of the strains sequenced here are not closely related to *S.* Typhimurium lineages circulating among cattle and humans in the United States. Below, we discuss some of these major African *S.* Typhimurium clades in detail (see the Supplemental Text for discussions of additional lineages).

**Figure 7.**
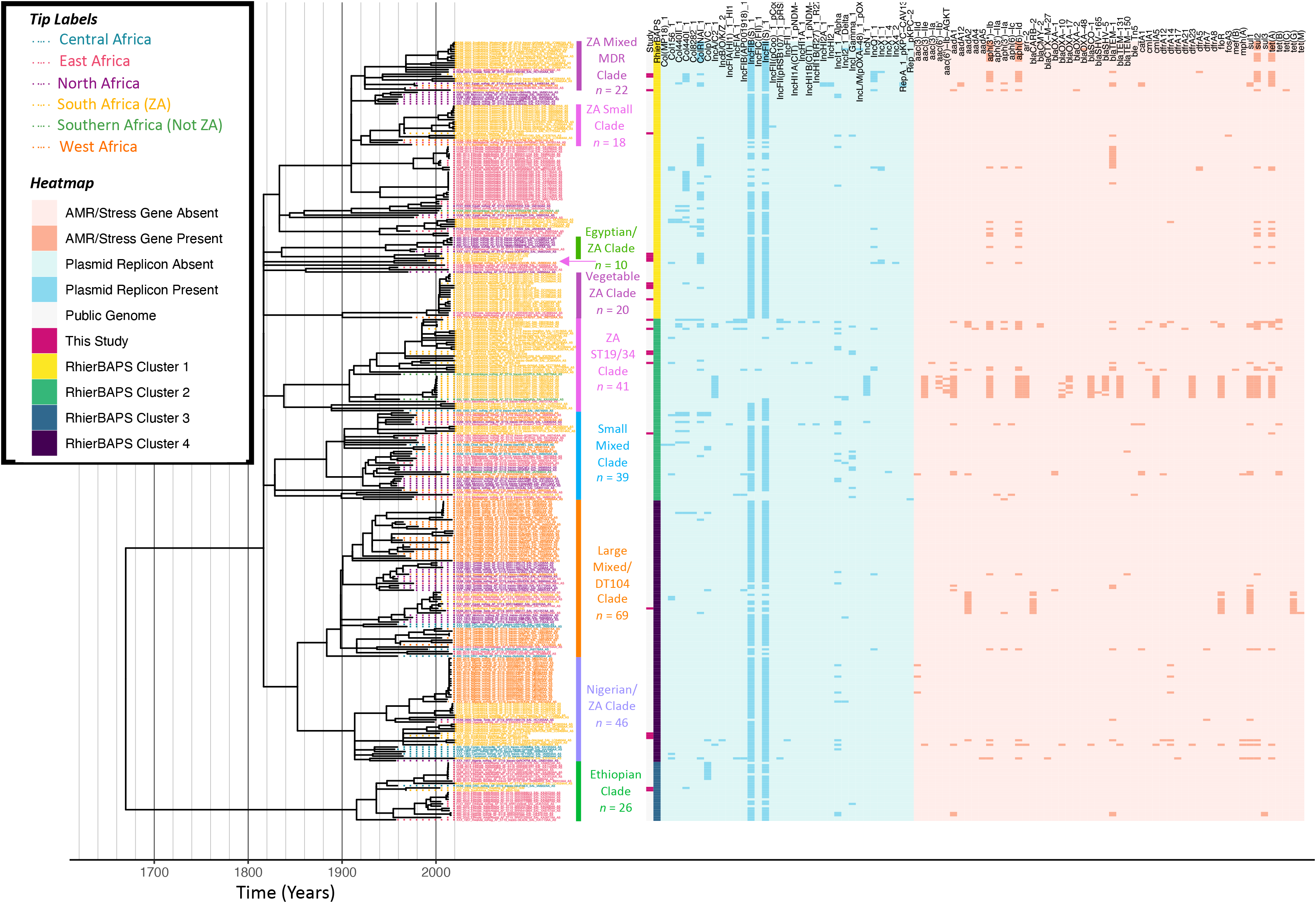
Maximum likelihood phylogeny constructed using core SNPs identified among 343 African *S.* Typhimurium genomes (319 publicly available genomes, plus the 24 sequenced here). Tip label colors denote the region/country from which each strain was reported to have been isolated (based on African regions as defined by the African Union, 25 April 2021). Clade labels denote clades discussed in either the main manuscript or the Supplemental Text. The heatmap to the right of the phylogeny denotes: (i) whether an isolate was sequenced in conjunction with this study (dark pink) or not (gray; “Study”); (ii) level 1 cluster assignments obtained using RhierBAPS (“RhierBAPS”); the presence and absence of (iii) plasmid replicons (blue) and (iv) antimicrobial resistance (AMR) determinants (orange). The phylogeny was rooted and time-scaled using LSD2, with branch lengths reported in years (X-axis). Core SNPs were identified among all genomes using Parsnp. AMR determinants were identified using ABRicate, the NCBI AMR determinant database, and minimum identity and coverage thresholds of 75 and 50%, respectively. Plasmid replicons were identified using ABRicate and the PlasmidFinder database, using minimum identity and coverage thresholds of 80 and 60%, respectively. The phylogeny was constructed and annotated using IQ-TREE and bactaxR/ggtree, respectively.

### A *S.* Typhimurium DT104-like clade emerged in Africa in the twentieth century as antimicrobial-susceptible and later acquired MDR

A *S.* Typhimurium strain sequenced here (PIG_2002_FS_040ST-S45), isolated in 2002 from swine meat in South Africa’s Free State province, clustered within a 69-isolate clade, which shared a common ancestor dated circa 1884 (95% CI [1153.59, 1956.00]; denoted as the “Large Mixed/DT104 Clade” in Figure 7). This large clade contained a mixture of human-, animal-, environmental, and food-associated isolates from Senegal (*n* = 16), Gambia (*n* = 13), Tunisia (*n* = 10), Benin (*n* = 7), Ethiopia (*n* = 6), Morocco (*n* = 4), DRC and South Africa (*n* = 3 each), Madagascar (*n* = 2), Algeria, Cameroon, Egypt, Kenya, and Tanzania (*n* = 1 each). The strain isolated in this study possessed five AMR genes: streptomycin resistance-conferring *aadA2*, beta lactamase *bla_CARB-2_,* chloramphenicol resistance-conferring *floR*, sulfonamide resistance-conferring *sul1*, and tetracycline resistance-conferring *tet(G)* (Figure 7). Notably, when compared to genomes from a previous study of *S.* Typhimurium in the United States (New York State) (32), this isolate clustered among DT104 strains isolated from dairy cattle and humans, sharing a common ancestor dated circa 1975 (95% CI [1467.42, 1999.00]; Supplemental Figure S6).

Within the Large Mixed/DT104 Clade in the African *S.* Typhimurium phylogeny (Figure 7), the DT104-like strain sequenced here (PIG_2002_FS_040ST-S45) was part of a 14-isolate subclade, which shared a common ancestor dated circa 1939 (95% CI [1623.18, 1960.00]). Notably, the four most distant members of this subclade, corresponding to strains isolated (i) in 1960 from a dog in Algeria, (ii) in 1970 and (iii) 1975 from unknown sources in Morocco, and (iv) in 1967 from a human in Morocco, were the only strains within this subclade that did not possess any AMR genes (Figure 7). The Moroccan strain isolated in 1975 was additionally reported to have itself been phage typed as DT104. The remaining ten genomes, which included the DT104-like strain sequenced here, clustered together, sharing a common ancestor dated circa 1980 (95% CI [1932.92, 2001.00]; Figure 7). Nine of these ten genomes possessed the five AMR genes listed above, which confer resistance to ampicillin, chloramphenicol, streptomycin, sulfonamides, and tetracycline (ACSSuT; one 2005 isolate from a camel in Ethiopia clustered among these genomes, but possessed only *aadA2* and *sul1*, while another strain, isolated in 2005 from poultry in Ethiopia, possessed all five AMR genes, as well as kanamycin resistance gene *aph(3’)-Ia* and tetracycline resistance gene *tet(M)*; Figure 7). This is noteworthy, as the ACSSuT AMR profile is often seen as characteristic of MDR D104 (33). Using the most parsimonious explanation for the acquisition of its MDR phenotype, the clade of African DT104-like isolates identified here emerged as antimicrobial-susceptible circa 1939 (95% CI [1623.18, 1960.00]) and acquired the MDR phenotype between ≍1966 and ≍1980 (95% CI [1820.78, 2001.00]; Figure 7).

### An antimicrobial-susceptible *S.* Typhimurium clade, which emerged in South Africa after 2000, encompasses isolates from produce, fish, poultry, and avian sources

Four additional isolates sequenced in this study were contained within a 20-isolate clade, which shared a common ancestor dated to circa 1900 (95% CI [1220.88, 1974.00]; denoted in Figure 7 as the “Vegetable ZA Clade”). No AMR genes were detected in any genomes within this clade, including the four strains sequenced here, which were all isolated from South Africa’s Western Cape province (two strains isolated in 2019 from fish food products, one in 2004 from ostrich feces, and one in 2003 from poultry meat; Figure 7). Interestingly, these isolates clustered among (i) 13 publicly available genomes, all derived from food-associated strains isolated in 2015 in South Africa (i.e., 4 from cabbage, 3 from carrots, 2 from lettuce, 2 from plant salad, and one from each of spinach and red onion) and (ii) one strain isolated from human feces in Addis Ababa, Ethiopia, sharing a common ancestor dated circa 2003 (95% CI [1746.59, 2003.00]; Figure 7). Members of this clade additionally shared a common ancestor with a strain isolated in 2013 from swine feces in Addis Ababa, Ethiopia, which was predicted to have existed circa 1990 (95% CI [1620.12, 2003.00]; Figure 7). The most distant member within the clade was a strain isolated in 1974 from Burkina Faso (Figure 7).

## DISCUSSION

### Endemic *Salmonella enterica* lineages are circulating among animals and animal products in South Africa and may infect humans

Geography plays an important role in shaping bacterial pathogen population structure, including that of *Salmonella enterica*, and different geographic regions may harbor their own endemic lineages (31, 34–38). Here, we observed numerous endemic *Salmonella enterica* lineages circulating among animals and animal products in South Africa, some of which encompassed human clinical isolates. Within the global *S.* Dublin ST10 phylogeny, for example, we identified a largely AMR-susceptible South Africa-specific clade, which has been circulating among animals, foods, and humans in the country for decades. While *S.* Dublin is largely considered to be a bovine-adapted serotype, human infections caused by *S.* Dublin are frequently invasive and may result in severe illness and/or death (5-11, 39, 40. In South Africa, invasive non-typhoidal salmonellosis is a serious public health concern: in 2019, over 25% of all non-typhoidal salmonellosis cases reported to the Group for Enteric, Respiratory and Meningeal disease Surveillance in South Africa (GERMS-SA) were invasive (825 and 2,437 reported invasive and non-invasive non-typhoidal salmonellosis cases, respectively; 25.3%) (41). While *S.* Dublin is not among the most common serotypes isolated from human clinical cases in South Africa (41), routine veterinary surveillance has revealed that *S.* Dublin is frequently isolated from animal sources in the country, particularly cattle (42). Further WGS efforts are needed to provide insight into the evolution and between-host transmission dynamics of the endemic South African *S.* Dublin lineage identified in this study.

In addition to the endemic South African *S.* Dublin lineage, we identified a clade of South African animal- and animal product-associated *S.* Hadar ST33 strains, which clustered near the root of the global *S.* Hadar ST33 phylogeny. First described as a novel *Salmonella* serotype in 1954 (43), *S.* Hadar was reported to have been responsible for several cases of diarrheal illness in Israel (43). Reportedly, serotype *S.* Hadar was rarely isolated prior to 1971; however, in the mid-1970s, *S.* Hadar quickly became the second-most common cause of human nontyphoidal salmonellosis in the United Kingdom (44–46). Consistent with these observations, the South African *S.* Hadar clade identified here shared a common ancestor dated circa 1962 and contained strains isolated from the 1960s through 2018. In South Africa specifically, *S.* Hadar is not among the top serotypes associated with human clinical cases (41); however, *S.* Hadar has been commonly isolated from animals and animal associated-environments in the country for nearly two decades (42, 47, 48). The results presented here indicate that an endemic *S.* Hadar ST33 lineage has been circulating among animals in South Africa for over fifty years; however, future WGS efforts querying *S.* Hadar strains from around the world—historical strains isolated prior to the 1970s, in particular—are needed to refine estimates as to when this particular lineage emerged.

In addition to the *S.* Dublin ST10 and *S.* Hadar ST33 endemic South African lineages, we observed that African *S.* Enteritidis ST11 could largely be partitioned into one largely antimicrobial-susceptible and one largely MDR clade. South African *S.* Enteritidis ST11, including those sequenced here, were confined to the largely antimicrobial-susceptible clade. These results are consistent with those observed in a previous study of *S.* Enteritidis in Africa (25), in which a geographically distinct MDR *S.* Enteritidis lineage was identified in Africa’s Central/East regions and rarely detected in South Africa. Since 2012, *S.* Enteritidis has been the serotype most commonly isolated from human clinical cases in South Africa (41). Among animals, *S.* Enteritidis has been one of the most frequently isolated serotypes in South Africa for decades, particularly from poultry-associated sources (42, 47, 48). Our results further support that South African *S.* Enteritidis, which is one of the most common *Salmonella enterica* serotypes circulating among animals and humans in the country, acquires AMR only sporadically and is, on a genomic scale, distinct from MDR *S.* Enteritidis lineages circulating in other regions of Africa. Collectively, our study reveals that endemic lineages of several non-typhoidal *Salmonella enterica* serotypes are circulating among animals and animal products in South Africa, some of which may occasionally infect humans.

### WGS can differentiate endemic and ecdemic *Salmonella enterica* lineages

Pathogenic bacteria not previously endemic to a given geographic region can be introduced into that region through the movement of humans, food, and/or animals (23, 36, 49, 50). In addition to observing *Salmonella enterica* lineages that were likely endemic to South Africa, our study identified numerous lineages that were likely to have been introduced into the country only recently. One *S.* Dublin isolate sequenced in this study, for example, clustered among isolates from the United States, indicating that this strain had been introduced into South Africa only recently. *S.* Dublin from the United States has previously been shown to be distinct from *S.* Dublin strains isolated in other world regions on a genomic scale (31), and the United States was one of the leading poultry exporters to South Africa in 2016 (i.e., the year the ecdemic *S.* Dublin strain sequenced here was isolated) (51). Our recent introduction hypothesis was further supported by metadata indicating that this strain had been isolated from poultry meat imported from North America and sold in a supermarket in South Africa’s Eastern Cape province.

We observed similar results for *S.* Hadar: one *S.* Hadar strain sequenced in this study was more closely related to *S.* Hadar from the United States than to its South African counterparts, which all formed a clade near the global *S.* Hadar ST33 phylogeny root (44–46). Unlike the *S.* Dublin strain sequenced here, which was likely introduced into South Africa from imported poultry meat, it is unclear exactly how the unique *S.* Hadar lineage sequenced here was introduced into the country, as its representative strain was isolated from bovine feces in 1990 and shared a common ancestor circa 1982 with a canine-associated *S.* Hadar strain isolated in 1988 in the United States. Future WGS efforts querying *S.* Hadar may provide insight into this lineage and its emergence in South Africa.

The *S.* Typhimurium isolates sequenced here were distributed across the African *S.* Typhimurium phylogeny, indicating that South Africa harbors numerous *S.* Typhimurium lineages. Since 2012, *S.* Typhimurium has been the second-most common non-typhoidal *Salmonella enterica* serotype isolated from human clinical cases in South Africa (after *S.* Enteritidis) and in 2019 was the most common serotype isolated from human clinical cases in the Eastern Cape province (41). *S.* Typhimurium has additionally been one of the most frequently isolated serotypes from animals and wildlife in South Africa for decades, and it is frequently isolated from a broad range of hosts (e.g., cattle, poultry, equine, sheep/goats, feline, rhinoceros) (42, 47, 48). Interestingly, we identified a largely AMR-susceptible, primarily South African *S.* Typhimurium clade, which contained isolates from produce, fish, poultry, and avian sources, and one human clinical isolate from Ethiopia (referred to above as the “Vegetable ZA Clade”), which was predicted to have been introduced into the country recently (i.e., after the year 2000). It is unclear exactly where this lineage originated and how it was introduced into South Africa, but future WGS efforts may elucidate this.

We additionally identified a *S.* Typhimurium clade, which contained the genomes of strains assigned to phage type DT104. MDR DT104 was responsible for a global epidemic in the 1990s, during which it was increasingly isolated from a broad range of animals (e.g., cattle, poultry, pigs, sheep), as well as human clinical cases (52). Notably, DT104 was predicted to have emerged as antimicrobial-susceptible circa 1948, later acquiring its MDR phenotype circa 1972 (33). The results observed here are consistent with these findings, as the DT104 clade identified here emerged as antimicrobial-susceptible circa 1939 and acquired the MDR phenotype between ≍1966 and ≍1980. The DT104 clade identified here spanned multiple African regions, and South African DT104-like genomes were distributed across the clade, indicating that South Africa may have been subjected to multiple DT104 introduction events and/or between-country transmission events; however, the lack of available DT104-like genomes from South Africa (i.e., one sequenced here and two publicly available genomes) and the African continent as a whole limits our ability to say this conclusively. Taken together, our results further highlight the strengths of WGS in *Salmonella* source tracking, both within and between countries and continents (53–55), and showcase the ability of WGS-based approaches to differentiate endemic and ecdemic lineages.

### WGS of historical isolates from under-sequenced geographic regions can provide novel insights into pathogen evolution and diversity

Worldwide, *Salmonella enterica* has been estimated to be responsible for more than 93 million illnesses and more than 150,000 deaths annually (56). In Africa, the disease burden imposed by *Salmonella enterica* is particularly significant; mortality and disability adjusted life years (DALYs) due to diarrheal disease and invasive infections caused by non-typhoidal serotypes are consistently higher in Africa than in other world regions (57). However, despite the disproportionally high incidence and burden of salmonellosis and other foodborne illnesses, the bulk of publicly available genomic data derived from *Salmonella enterica* has come from regions with lower burdens (57, 58); for example, among all *Salmonella enterica* genomes in Enterobase (accessed 7 April 2021), over 80% were derived from strains reported to have been isolated in North America and Europe (128,517 and 104,910 genomes from North America and Europe, respectively; 233,427 of 291,362 total *Salmonella enterica* genomes).

Here, we used WGS to characterize 63 *Salmonella enterica* strains isolated from animals and animal products in South Africa over a 60-year time span, which, to our knowledge, represents the most extensive WGS-based characterization of non-human-associated non-typhoidal *Salmonella enterica* in the country to date. Importantly, numerous genomes sequenced here belonged to lineages that were phylogenetically distinct from those circulating in more heavily sequenced/sampled regions of the world, such as North America and Europe. For example, as observed here, some African *S.* Dublin ST10 isolates do not belong to the two major *S.* Dublin ST10 clades circulating primarily in North America and Europe, indicating that *S.* Dublin isolates representing clades outside of the two major North American- and European-associated clades are likely circulating in other countries around the world, including African countries outside of South Africa. Similarly, the few available *S.* Enteritidis ST366 genomes are derived from strains primarily isolated in Africa. Future WGS efforts in Africa will likely provide insight into the evolution and emergence of these lineages, as well as novel clades and those underrepresented in public databases. Overall, this study offers a glimpse into the genomics of non-typhoidal *Salmonella enterica* lineages circulating among livestock, domestic animals, wildlife, and animal products in South Africa. Future WGS-based studies querying greater numbers of isolates from animal, food, and environmental sources are needed to better understand the evolution, population structure, and AMR dynamics of this important pathogen.

## MATERIALS AND METHODS

### Isolate selection

The isolates used in this study were recovered from samples submitted between 1957 and 2019 at Bacteriology laboratory: Onderstepoort Veterinary Research, South Africa, as part of routine diagnostics services which includes isolation and serotyping of *Salmonella* strains. Therefore, a total of 73 isolates representing (i) four major *Salmonella enterica* serotypes (i.e., Dublin, Enteritidis, Hadar, and Typhimurium) in the country (42, 48) from (ii) various geographical locations in the country, (iii) different sources of isolation (animal and animal products), and (iv) animal species (livestock, companion animals, wildlife) were randomly selected for sequencing in this study. The isolates were preserved as lyophilized and revived by inoculation into brain heart infusion (BHI) broth and incubated at 37°C for 18-24 hours.

### Whole-genome sequencing

Genomic DNA was extracted from BHI broth cultures using the High Pure PCR template preparation kit (Roche, Potsdam, Germany) according to the manufacturer’s instructions. WGS of the isolates was performed at the Biotechnology Platform, Agricultural Research Council, South Africa. DNA libraries were prepared using TruSeq and Nextera DNA library preparation kits (Illumina, San Diego, CA, USA), followed by sequencing on Illumina HiSeq and MiSeq instruments (Illumina, San Diego, CA, USA).

### Initial data processing and quality control

Quality control, adapter removal, decontamination, and error correction of the raw sequencing data was performed using BBDuk v. 37.90 (https://jgi.doe.gov/data-and-tools/ bbtools/bb-tools-user-guide/bbduk-guide/), and SPAdes v. 3.12.0 (59) was used to create a *de novo* assembly for each isolate. FastQC v. 0.11.5 (https://www.bioinformatics.babraham.ac.uk/projects/fastqc/) was used to assess the quality of the paired-end reads associated with each isolate (*n* = 73 isolates total; 21, 15, 11, and 26 isolates assigned to serotypes Dublin, Enteritidis, Hadar, and Typhimurium, respectively) (60), and QUAST v. 4.5 (61) was used to assess the quality of the associated assembled genome (Supplemental Table S1). The lineage workflow (i.e., “lineage_wf”) implemented in CheckM v. 1.1.3 (62) was additionally used to identify potential contamination in each assembled genome, as well as to assess genome completeness (Supplemental Table S1). MultiQC v. 1.8 (63) was used to assess the quality of all genomes in aggregate. Several low-quality isolate genomes with >5% contamination and/or <95% completeness were identified (*n* = 3, 2, 3, and 2 low-quality isolate genomes assigned to serotypes Dublin, Enteritidis, Hadar, and Typhimurium, respectively) and were thus omitted from further analysis, yielding a final set of 63 *Salmonella enterica* genomes used in subsequent steps (Supplemental Table S1).

### *In silico* serotyping and multi-locus sequence typing

All 63 assembled *Salmonella enterica* genomes (see section “Initial data processing and quality control” above) underwent *in silico* serotyping using the command line implementations of (i) the *Salmonella In Silico* Typing Resource (SISTR) v. 1.1.1 (18) and (ii) the *k*-mer based workflow implemented in SeqSero2 v. 1.1.1 (17) (Supplemental Table S1). Each genome additionally underwent *in silico* seven-gene MLST using mlst v. 2.9 (https://github.com/tseemann/mlst) and the seven-gene scheme available for *Salmonella enterica* (--scheme ‘senterica’) in PubMLST (64, 65) (Supplemental Table S1).

### Reference-free SNP identification and phylogeny construction

The 63 *Salmonella enterica* genomes sequenced in this study were compared to 442 of the 445 *Salmonella* genomes described by Worley, et al. (28) (three genomes did not have publicly available sequence read archive [SRA] data at the time of access, i.e., 20 February 2019). The SRA toolkit v. 2.9.6 was used to download paired-end reads for each of the 442 publicly available genomes (66, 67), which were then assembled into contigs using SPAdes v. 3.8.0 (59), using *k*-mer sizes of 21, 33, 55, 77, 99, and 127, and the “careful” option. SNPs were identified among all 505 assembled *Salmonella* genomes with kSNP3 v. 3.92 (68, 69), using the optimal *k*-mer size determined by Kchooser (*k* = 19). The resulting core SNP alignment was supplied as input to IQ-TREE v. 1.5.4 (70), which was used to construct a ML phylogeny using the optimal ascertainment bias-aware nucleotide substitution model identified using ModelFinder (based on its Bayesian Information Criteria [BIC] value) (71) and 1,000 replicates of the Ultrafast Bootstrap method (72, 73). The resulting ML phylogeny (Figure 1) was annotated using FigTree v. 1.4.4 (http://tree.bio.ed.ac.uk/software/figtree/). All reference-free SNP identification and ML phylogeny construction steps described above were repeated to identify SNPs among the 63 *Salmonella enterica* genomes sequenced here, with publicly available genomes excluded; the resulting ML phylogeny was annotated in R v. 3.6.1 (74) using the bactaxR package (75) and its dependencies ggtree (76, 77), ape (78), dplyr (79), phylobase (80), phytools (81), and reshape2 (82) (Figure 2 and Supplemental Table S2).

### *In silico* AMR determinant, plasmid replicon, and virulence factor detection

AMR determinants were identified within each of the 63 *Salmonella* genomes sequenced in this study, using each of the following pipelines: (i) AMRFinderPlus v. 3.9.3 (29), (ii) ABRicate v. 1.0.1 (https://github.com/tseemann/abricate), and (iii) ARIBA v. 2.14.6 (83) (Figure 2 and Supplemental Figure S1). For the AMRFinderPlus pipeline, Prokka v. 1.13 (84) was used to annotate each of the 63 assembled genomes; the resulting GFF (.gff) and FASTA (.faa and .ffn) files were used as input for AMRFinderPlus, which was used to identify AMR and stress response determinants in each genome, using the *Salmonella* organism option and the most recent AMRFinderPlus database (database v. 2020-11-09.1, accessed 21 November 2020). For the ABRicate pipeline, AMR determinants were identified in each assembled genome using the NCBI AMR database (--db ncbi; accessed 19 April 2020) (29) and minimum identity and coverage thresholds of 75 (--minid 75) and 50% (--mincov 50), respectively. For the ARIBA pipeline, ARIBA’s getref and prepareref commands were used to download and prepare the latest version of the ResFinder database (accessed 14 February 2021), respectively (85). ARIBA’s run command was then used to identify AMR determinants in each genome, using the paired-end reads associated with each isolate as input.

ABRicate and ARIBA were additionally used to detect plasmid replicons within each of the 63 *Salmonella* genomes sequenced in this study using the PlasmidFinder database (30) (Figure 2 and Supplemental Figure S2). For the ABRicate pipeline, assembled genomes were used as input, and plasmid replicons were detected in each genome (--db plasmidfinder; PlasmidFinder database accessed 19 April 2020) using minimum identity and coverage thresholds of 80 (--minid 80) and 60% (--mincov 60), respectively. For the ARIBA pipeline, ARIBA’s getref and prepareref commands were used to download and prepare the latest version of the PlasmidFinder database (accessed 14 February 2021), respectively. ARIBA’s run command was then used to identify plasmid replicons in each genome, using paired-end reads associated with each isolate as input. ABRicate was further used to detect virulence factors in each genome, using the Virulence Factor Database (VFDB; --db vfdb, accessed 19 April 2020) (86, 87), using minimum identity and coverage thresholds of 70 (--minid 70) and 50% (--mincov 50), respectively (Figure 2 and Supplemental Table S2).

### Construction of time-scaled *S.* Dublin phylogenies

To compare the 18 *S.* Dublin isolates sequenced in this study to publicly available *S.* Dublin genomes, all genomes meeting each of the following conditions were downloaded via Enterobase (accessed 27 December 2020, *n* = 2,784; Supplemental Table S3): (i) genomes were assigned to sequence type (ST) 10 (i.e., the ST to which all of the *S.* Dublin isolates sequenced in this study were assigned/approximately assigned) using the Achtman seven-gene MLST scheme for *Salmonella*; (ii) genomes had an exact year of isolation reported in Enterobase’s “Collection Year” field; (iii) genomes could be assigned to a known isolation source, with “Laboratory” strains excluded, per Enterobase’s “Source Niche” field; (iv) genomes could be assigned to a known country of isolation, per Enterobase’s “Country” field (88, 89). All 2,802 assembled *S.* Dublin genomes underwent *in silico* plasmid replicon and AMR determinant detection using ABRicate v. 1.0.1 and the PlasmidFinder and NCBI AMR databases, respectively, as described above (see section “*In silico* AMR determinant, plasmid replicon, and virulence factor detection” above).

Parsnp and HarvestTools v. 1.2 (90) were used to identify core SNPs among all 2,802 *S.* Dublin genomes (2,784 publicly available genomes, plus the 18 sequenced here), using the closed chromosome of ST10 *S.* Dublin str. USMARC-69838 (NCBI Nucleotide Accession NZ_CP032449.1) as a reference and Parsnp’s implementation of PhiPack to remove recombination (91). Clusters were identified within the resulting core SNP alignment using RhierBAPs v. 1.1.3 (92, 93), R v. 4.0.0, and three clustering levels. IQ-TREE v. 1.5.4 (70) was used to construct a ML phylogeny using (i) the resulting core SNPs as input; (ii) an ascertainment bias correction (to account for the use of solely variant sites), corresponding to constant sites estimated using the GC content of the reference chromosome (-fconst 1171365,1282543,1281883,117722); (iii) the optimal nucleotide substitution model selected using ModelFinder (71), based on its corresponding BIC value (i.e., the TVM+I model); (iv) 1,000 replicates of the UltraFast bootstrap approximation (72).

The resulting ML phylogeny was rooted and time-scaled using LSD2 v. 1.4.2.2 (94) and the following parameters: (i) tip dates corresponding to the year of isolation associated with each genome; (ii) a fixed substitution rate of 2.79×10^-7^ substitutions/site/year (i.e., the substitution rate estimated in a previous study of *S.* Typhimurium phage type DT104) (33); (iii) constrained mode (-c), with the root estimated using constraints on all branches (-r as); (iv) variances calculated using input branch lengths (-v 1); (v) 1,000 samples for calculating confidence intervals for estimated dates (-f 1000); (vi) a sequence length of 4,913,018 (i.e., the length of the reference chromosome; -s 4913018). The resulting phylogeny was annotated using the bactaxR package in R (Figure 3). All aforementioned *S.* Dublin SNP calling and phylogeny construction steps were repeated to construct time-scaled ML phylogenies using the following subsets of *S.* Dublin genomes: (i) members of a large *S.* Dublin clade, which contained all 18 *S.* Dublin isolates sequenced in this study (i.e., “*S.* Dublin Major Clade I”, *n* = 1,787 genomes; Supplemental Figure S3); (ii) a smaller clade within *S.* Dublin Major Clade I, which contained 17 of the 18 *S.* Dublin isolates sequenced here (i.e., the “*S.* Dublin Small Subclade”, *n* = 78; Figure 4); (iii) a larger clade within *S.* Dublin Major Clade I, which contained one *S.* Dublin isolate sequenced here (i.e., the “*S.* Dublin Large Subclade”, *n* = 1,709; Supplemental Figure S4).

### Construction of time-scaled *S.* Hadar phylogeny

To compare the eight *S.* Hadar isolates sequenced in this study to publicly available *S.* Hadar genomes, all genomes meeting each of the following conditions were downloaded via Enterobase (accessed 10 January 2021, *n* = 1,562; Supplemental Table S4): (i) genomes were assigned to ST33 (i.e., the ST to which all of the *S.* Hadar isolates sequenced in this study were assigned) using the Achtman seven-gene MLST scheme for *Salmonella*; (ii) genomes had an exact year of isolation reported in Enterobase’s “Collection Year” field; (iii) genomes could be assigned to a known isolation source, with “Laboratory” strains excluded, per Enterobase’s “Source Niche” field; (iv) genomes could be assigned to a known country of isolation, per Enterobase’s “Country” field (88, 89). All 1,570 assembled *S.* Hadar genomes underwent *in silico* plasmid replicon and AMR determinant detection using ABRicate v. 1.0.1 and the PlasmidFinder and NCBI AMR databases, respectively, as described above (see section “*In silico* AMR determinant, plasmid replicon, and virulence factor detection” above).

Parsnp and HarvestTools v. 1.2 (90) were used to identify core SNPs among all 1,570 *S.* Hadar genomes (1,562 publicly available genomes, plus the eight sequenced here), using the closed chromosome of ST33 *S.* Hadar str. FDAARGOS_313 (NCBI Nucleotide Accession NZ_CP022069.2) as a reference and Parsnp’s implementation of PhiPack to remove recombination (91). Clusters were identified within the resulting core SNP alignment using RhierBAPs v. 1.1.3 (92, 93), R v. 4.0.0, and three clustering levels. IQ-TREE v. 1.5.4 (70) was used to construct a ML phylogeny using (i) the resulting core SNPs as input; (ii) an ascertainment bias correction (to account for the use of solely variant sites), corresponding to constant sites estimated using the GC content of the reference chromosome (-fconst 1179063,1283051,1279961,1174705); (iii) the optimal nucleotide substitution model selected using ModelFinder (71), based on its corresponding BIC value (i.e., the K3Pu+I model) (95); (iv) 1,000 replicates of the UltraFast bootstrap approximation (72). All aforementioned SNP calling and phylogeny construction steps were repeated, with a single outlier genome from the United Kingdom (Enterobase Assembly Barcode SAL_GB0368AA_AS) removed, yielding a 1,569-isolate *S.* Hadar phylogeny that was used in subsequent steps.

The resulting ML phylogeny was rooted and time-scaled using LSD2 v. 1.4.2.2 (94) and the following parameters: (i) tip dates corresponding to the year of isolation associated with each genome; (ii) a fixed substitution rate of 2.79×10^-7^ substitutions/site/year (i.e., the substitution rate estimated in a previous study of *S.* Typhimurium phage type DT104) (33); (iii) constrained mode (-c), with the root estimated using constraints on all branches (-r as); (iv) variances calculated using input branch lengths (-v 1); (v) 1,000 samples for calculating confidence intervals for estimated dates (-f 1000); (vi) a sequence length of 4,916,780 (i.e., the length of the reference chromosome; -s 4916780). The resulting phylogeny was annotated using the bactaxR package in R (Figure 5).

### Construction of time-scaled *S.* Enteritidis phylogenies

To compare the 13 *S.* Enteritidis isolates sequenced in this study to publicly available *S.* Enteritidis genomes, all genomes meeting each of the following conditions were downloaded via Enterobase (accessed 27 December 2020, *n* = 697; Supplemental Table S5): (i) genomes were assigned to ST11 (i.e., the ST to which 12 of the 13 *S.* Enteritidis isolates sequenced in this study were assigned/approximately assigned) using the Achtman seven-gene MLST scheme for *Salmonella*; (ii) genomes had an exact year of isolation reported in Enterobase’s “Collection Year” field; (iii) genomes could be assigned to a known country of isolation within the African continent, per Enterobase’s “Country” and “Continent” fields, respectively (88, 89). Additionally, one isolate sequenced here was assigned to ST366, a ST that differs from ST11 by a single allele (i.e., *purE*). As such, all ST366 genomes available in Enterobase were additionally downloaded (*n =* 10), and those with known isolation years and country/continents of isolation (*n* = 6; three isolates from South Africa, two from Zambia, and one from the United Kingdom) were used in subsequent steps. All 716 assembled *S.* Enteritidis genomes underwent *in silico* plasmid replicon and AMR determinant detection using ABRicate v. 1.0.1 and the PlasmidFinder and NCBI AMR databases, respectively, as described above (see section “*In silico* AMR determinant, plasmid replicon, and virulence factor detection” above).

Parsnp and HarvestTools v. 1.2 (90) were used to identify core SNPs among all 716 *S.* Enteritidis genomes (703 publicly available genomes, plus the 13 sequenced here), using the closed chromosome of ST11 *S.* Enteritidis str. OLF-SE10-10052 (NCBI Nucleotide Accession NZ_CP009092.1) as a reference and Parsnp’s implementation of PhiPack to remove recombination (91). Clusters were identified within the resulting core SNP alignment using RhierBAPs v. 1.1.3 (92, 93), R v. 4.0.0, and three clustering levels. IQ-TREE v. 1.5.4 (70) was used to construct a ML phylogeny using (i) the resulting core SNPs as input; (ii) an ascertainment bias correction (to account for the use of solely variant sites), corresponding to constant sites estimated using the GC content of the reference chromosome (-fconst 1127671,1230753,1225740,1125726); (iii) the optimal nucleotide substitution model selected using ModelFinder (71), based on its corresponding BIC value (i.e., the TVM+I model); (iv) 1,000 replicates of the UltraFast bootstrap approximation (72).

The resulting ML phylogeny was rooted and time-scaled using LSD2 v. 1.4.2.2 (94) and the following parameters: (i) tip dates corresponding to the year of isolation associated with each genome; (ii) a fixed substitution rate of 2.20×10^-7^ substitutions/site/year (i.e., the substitution rate estimated in a previous study of *S.* Enteritidis) (96); (iii) constrained mode (-c), with the root estimated using constraints on all branches (-r as); (iv) variances calculated using input branch lengths (-v 1); (v) 1,000 samples for calculating confidence intervals for estimated dates (-f 1000); (vi) a sequence length of 4,709,890 (i.e., the length of the reference chromosome; -s 4709890). The resulting phylogeny was annotated using the bactaxR package in R (Figure 6). All aforementioned *S.* Enteritidis SNP calling and phylogeny construction steps were repeated to construct an additional time-scaled ML phylogeny using ST11 isolates within a major clade in the African *S.* Enteritidis phylogeny (referred to here as “African *S.* Enteritidis ST11 Major Clade I”, *n* = 517; Supplemental Figure S5). RhierBAPs v. 1.1.3 (92, 93) and R v. 4.0.0 were additionally used to identify clusters within the resulting core SNP alignment, using three clustering levels.

### Construction of time-scaled *S.* Typhimurium phylogenies

To compare the 24 *S.* Typhimurium isolates sequenced in this study to publicly available *S.* Typhimurium genomes, all genomes meeting each of the following conditions were downloaded via Enterobase (accessed 27 December 2020, *n* = 319; Supplemental Table S6): (i) genomes were assigned to either ST19 (the ST to which 23 of the 24 *S.* Typhimurium isolates sequenced in this study were assigned/approximately assigned) or ST34 (the ST of the remaining isolate, which differs from ST19 by a single allele, *dnaN*) using the Achtman seven-gene MLST scheme for *Salmonella*; (ii) genomes had an exact year of isolation reported in Enterobase’s “Collection Year” field; (iii) genomes could be assigned to a known country of isolation within the African continent, per Enterobase’s “Country” and “Continent” fields, respectively (88, 89). All 343 assembled *S.* Typhimurium genomes underwent *in silico* plasmid replicon and AMR determinant detection using ABRicate v. 1.0.1 and the PlasmidFinder and NCBI AMR databases, respectively, as described above (see section “*In silico* AMR determinant, plasmid replicon, and virulence factor detection” above).

Parsnp and HarvestTools v. 1.2 (90) were used to identify core SNPs among all 343 *S.* Typhimurium genomes (319 publicly available genomes, plus the 24 sequenced here), using the closed chromosome of ST19 *S.* Typhimurium str. LT2 (NCBI Nucleotide Accession NC_003197.2) as a reference and Parsnp’s implementation of PhiPack to remove recombination (91). Clusters were identified within the resulting core SNP alignment using RhierBAPs v. 1.1.3 (92, 93), R v. 4.0.0, and three clustering levels. IQ-TREE v. 1.5.4 (70) was used to construct a ML phylogeny using (i) the resulting core SNPs as input; (ii) an ascertainment bias correction (to account for the use of solely variant sites), corresponding to constant sites estimated using the GC content of the reference chromosome (-fconst 1160904,1268422,1268221,1159903); (iii) the optimal nucleotide substitution model selected using ModelFinder (71), based on its corresponding BIC value (i.e., the TVM+I model); (iv) 1,000 replicates of the UltraFast bootstrap approximation (72).

The resulting ML phylogeny was rooted and time-scaled using LSD2 v. 1.4.2.2 (94) and the following parameters: (i) tip dates corresponding to the year of isolation associated with each genome; (ii) a fixed substitution rate of 2.79×10^-7^ substitutions/site/year (i.e., the substitution rate estimated in a previous study of *S.* Typhimurium phage type DT104) (33); (iii) constrained mode (-c), with the root estimated using constraints on all branches (-r as); (iv) variances calculated using input branch lengths (-v 1); (v) 1,000 samples for calculating confidence intervals for estimated dates (-f 1000); (vi) a sequence length of 4,857,450 (i.e., the length of the reference chromosome; -s 4857450). The resulting phylogeny was annotated using the bactaxR package in R (Figure 7). All aforementioned *S.* Typhimurium SNP calling and phylogeny construction steps were repeated to construct an additional time-scaled ML phylogeny, using the 24 isolates sequenced here and 87 human- and bovine-associated *S.* Typhimurium isolates from a previous study of the serotype in New York State in the United States (32) (*n* = 111; Supplemental Figure S6).

### Data availability

Illumina reads for genomes sequenced in this study are available in the National Center for Biotechnology Information (NCBI) Sequence Read Archive (SRA) under BioProject accession PRJNA727588. Metadata for the *Salmonella enterica* genomes sequenced in this study are available in Supplemental Table S1. Enterobase (https://enterobase.warwick.ac.uk/) metadata for the publicly available genomes used in this study are available in Supplemental Tables S3-S6.

## ACKNOWLEDGMENTS

The following organizations and individuals are acknowledged for their contributions: (i) Gauteng Department of Agriculture and Rural Development (GDRAD) for funding this project; (ii) the officials from the Bacteriology section of ARC: OVR (Rosina Maluleka, Palesa Nthaba, Mmatua Motau and Lavhelesani Makhado) for the technical support during data retrieval; (iii) the authors are grateful to the Agricultural Research Council: Onderstepoort Veterinary Research for providing all research facilities. We also acknowledge our collaborators; EMBL.

