## Supplemental Figure S1 for "Genomic characterization of endemic and ecdemic non-typhoidal *Salmonella enterica* lineages circulating among animals and animal products in South Africa"

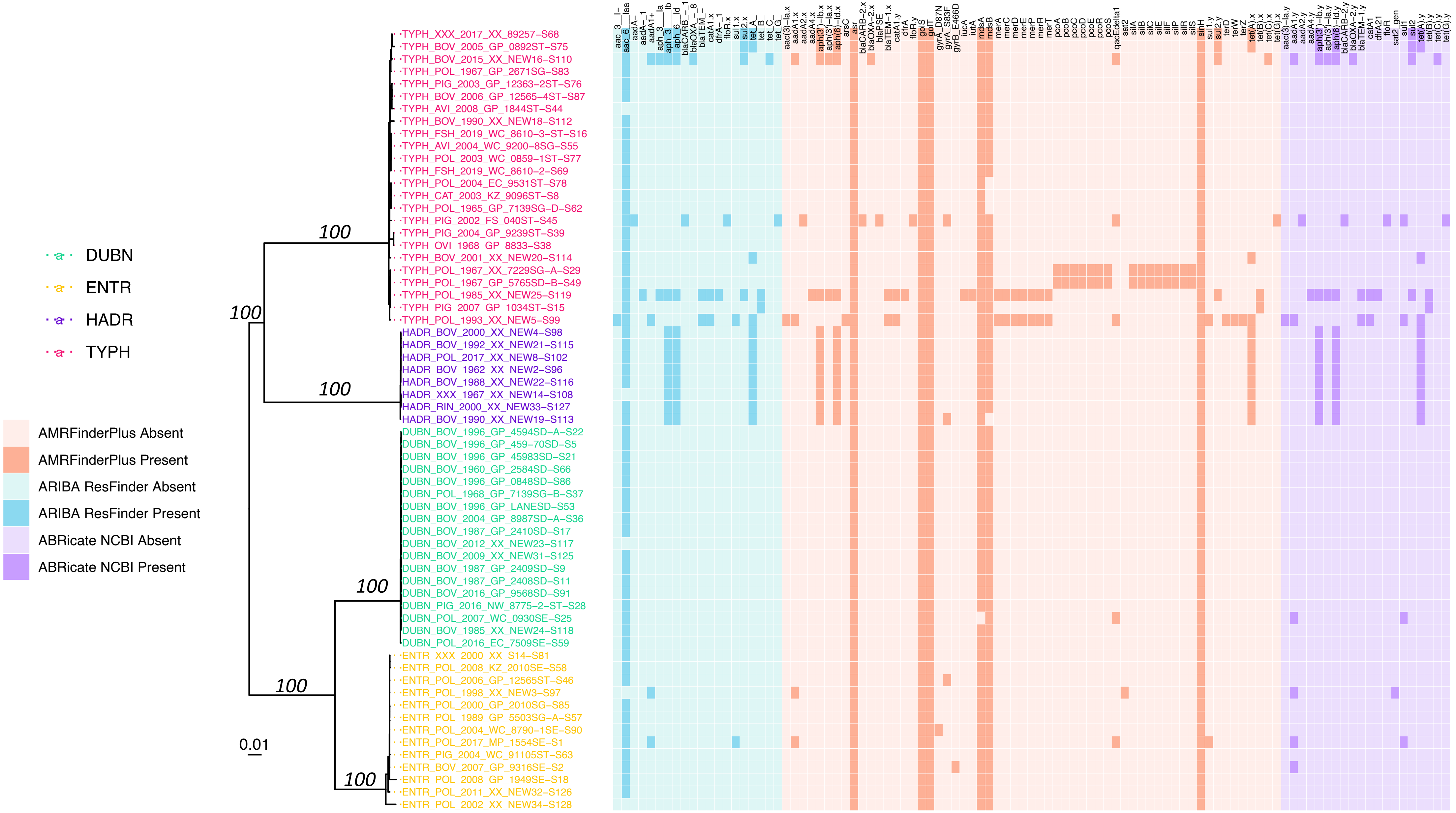


**Supplemental Figure S1.** Maximum likelihood phylogeny constructed using core SNPs identified among the genomes of 63 *Salmonella* strains isolated in conjunction with this study. Tip label colors denote isolate serotypes, and branch labels denote ultrafast bootstrap support percentages out of 1,000 replicates (selected for readability). The heatmap to the right of the phylogeny denotes the presence and absence of antimicrobial resistance (AMR) and/or stress response determinants detected in each genome using (i) ARIBA + the ResFinder AMR database (blue), (ii) AMRFinderPlus (orange), and (iii) ABRicate + the NCBI AMR database (purple). The phylogeny is rooted at the midpoint with branch lengths reported in substitutions per site. Core SNPs were identified among all genomes using kSNP3. AMR determinants identified using ABRicate were considered to be present in a genome at minimum identity and coverage thresholds of 75 and 50%, respectively. The phylogeny was constructed and annotated using IQ-TREE and bactaxR/ggtree, respectively.
