## Supplemental Figure S2 for "Genomic characterization of endemic and ecdemic non-typhoidal *Salmonella enterica* lineages circulating among animals and animal products in South Africa"

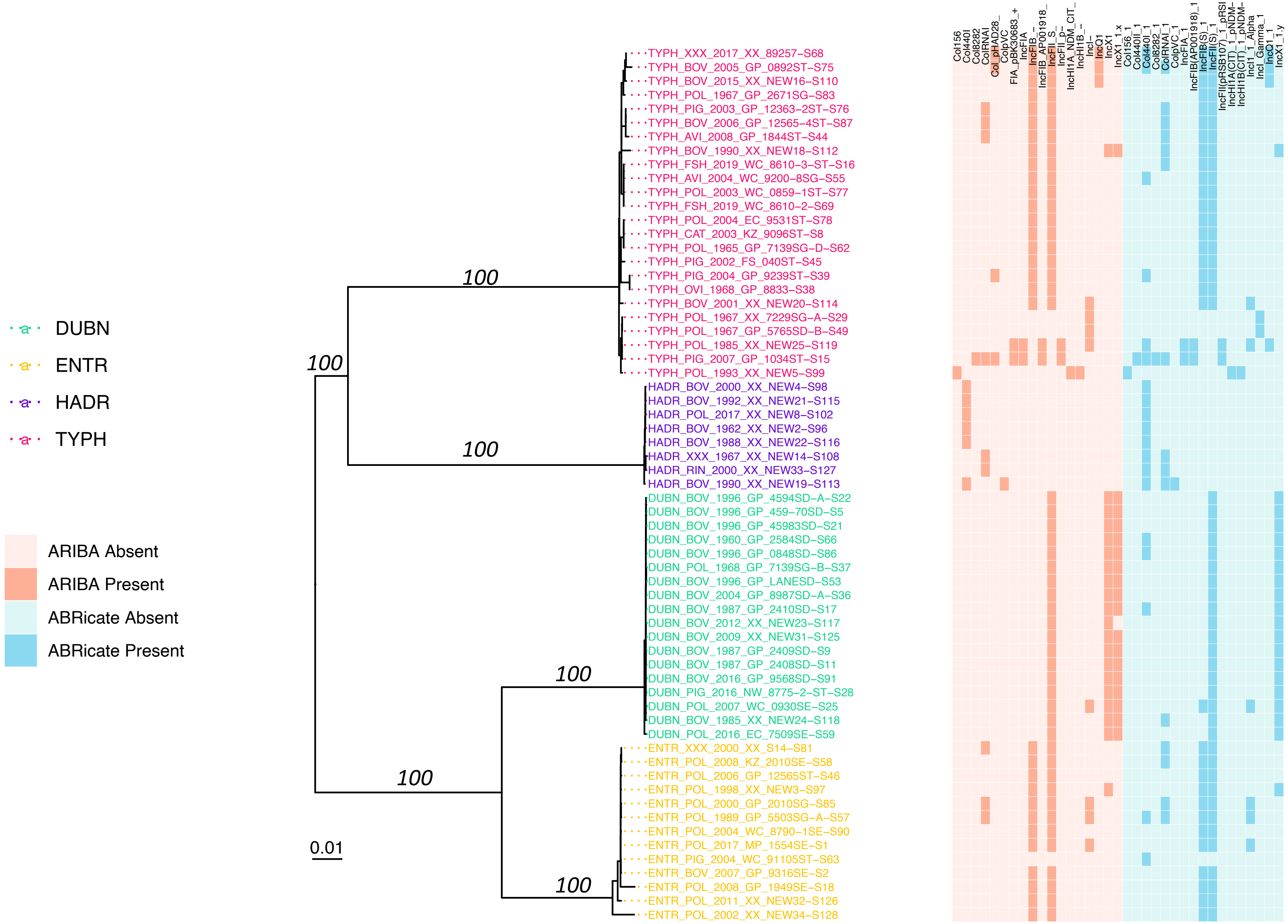


**Supplemental Figure S2.** Maximum likelihood phylogeny constructed using core SNPs identified among the genomes of 63 *Salmonella* strains isolated in conjunction with this study. Tip label colors denote isolate serotypes, and branch labels denote ultrafast bootstrap support percentages out of 1,000 replicates (selected for readability). The heatmap to the right of the phylogeny denotes the presence and absence of plasmid replicons detected in each genome using the PlasmidFinder database and (i) ARIBA (orange), and (ii) ABRicate (blue). The phylogeny is rooted at the midpoint with branch lengths reported in substitutions per site. Core SNPs were identified among all genomes using kSNP3. Plasmid replicons identified using ABRicate were considered to be present in a genome at minimum identity and coverage thresholds of 80 and 60%, respectively. The phylogeny was constructed and annotated using IQ-TREE and bactaxR/ggtree, respectively.
