## Supplemental Figure S4 for "Genomic characterization of endemic and ecdemic non-typhoidal *Salmonella enterica* lineages circulating among animals and animal products in South Africa"

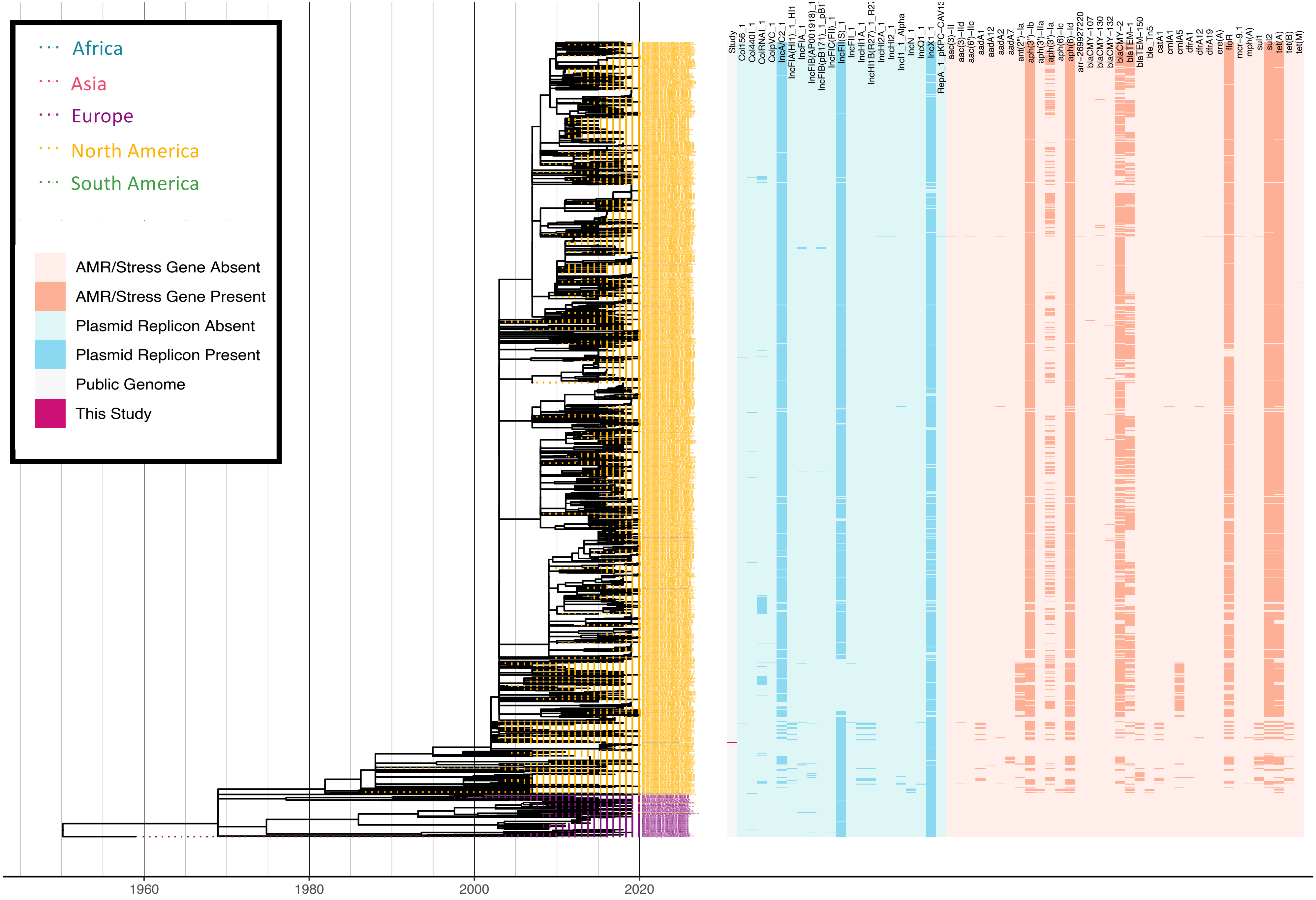


**Supplemental Figure S4.** Maximum likelihood phylogeny constructed using core SNPs identified among 1,709 *S.* Dublin genomes within the *S.* Dublin Large Subclade (1,708 publicly available genomes, plus one sequenced here). Tip label colors denote the continent from which each strain was reported to have been isolated. The heatmap to the right of the phylogeny denotes: (i) whether an isolate was sequenced in conjunction with this study (dark pink) or not (gray; "Study"); the presence and absence of (ii) plasmid replicons (blue) and (iii) antimicrobial resistance (AMR) determinants (orange). The phylogeny was rooted and time-scaled using LSD2, with branch lengths reported in years (X-axis). Core SNPs were identified among all genomes using Parsnp. AMR determinants were identified using ABRicate, the NCBI AMR determinant database, and minimum identity and coverage thresholds of 75 and 50%, respectively. Plasmid replicons were identified using ABRicate and the PlasmidFinder database, using minimum identity and coverage thresholds of 80 and 60%, respectively. The phylogeny was constructed and annotated using IQ-TREE and bactaxR/ggtree, respectively.
