## Supplemental Figure S6 for "Genomic characterization of endemic and ecdemic non-typhoidal *Salmonella enterica* lineages circulating among animals and animal products in South Africa"

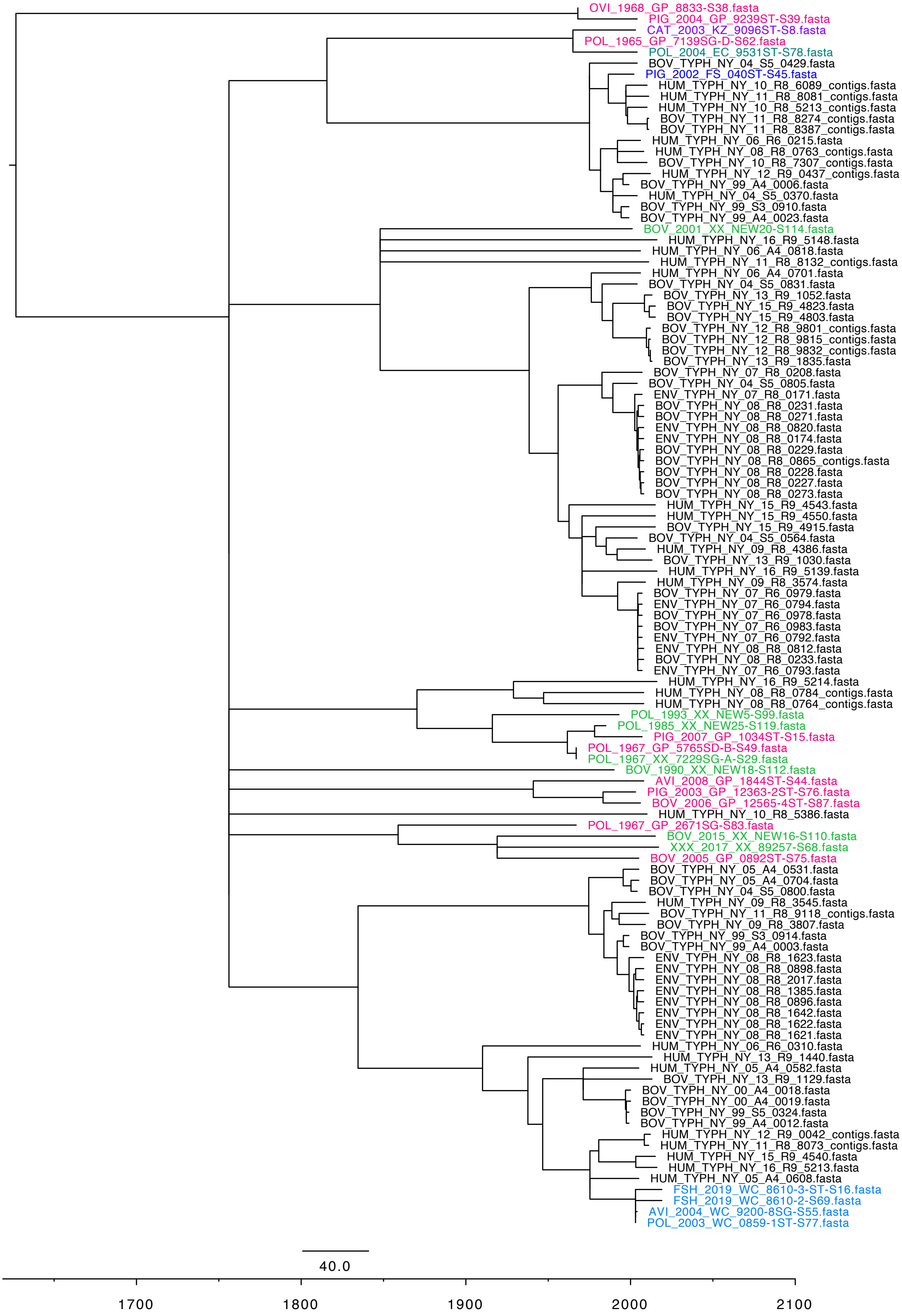


**Supplemental Figure S6.** Maximum likelihood phylogeny constructed using core SNPs identified among 111 *S.* Typhimurium genomes (87 publicly available genomes from a previous study of human- and bovine-associated *S.* Typhimurium in New York State, plus 24 sequenced here). Publicly available genomes from New York State (United States of America) are denoted by black tip labels, while genomes sequenced here are denoted by colored tip labels corresponding to the province from which their associated strains were isolated. The phylogeny was rooted and time-scaled using LSD2, with branch lengths reported in years (X-axis). Core SNPs were identified among all genomes using Parsnp. The phylogeny was constructed and annotated using IQ-TREE and FigTree, respectively.
