## Supplemental Text for "Genomic characterization of endemic and ecdemic non-typhoidal *Salmonella enterica* lineages circulating among animals and animal products in South Africa"

**Supplemental Text.** Descriptions of African *Salmonella enterica* serotype Typhimurium (*S.* Typhimurium) clades not discussed in the main manuscript.

**A primarily South African clade of ST19 and ST34 isolates contains a mixture of antimicrobial-susceptible and multidrug-resistant *S.* Typhimurium lineages.** Five isolates sequenced here were members of a 41-isolate clade, which shared a common ancestor dated circa 1838 (95% confidence interval [CI] of [1023.59, 1961.00]; denoted in Figure 7 as the "ZA ST19/34 Clade"). Among them was multidrug-resistant (MDR) sequence type 34 (ST34) strain POL_1993_XX_NEW5-S99, which was isolated from poultry meat in 1993 and possessed the following antimicrobial resistance (AMR) genes: gentamicin resistance gene *aac(3)-Ia,* streptomycin-resistance gene *aadA1*, beta lactamase $bla_{TEM-1}$, chloramphenicol-resistance gene *catA1*, sulfonamide resistance gene *sul1*, and tetracycline resistance gene *tet(A)* (Figure 7). This strain was part of a five-strain sub-clade, which contained all four publicly available African ST34 genomes, all of which were isolated from humans in South Africa; members of this ST34 sub-clade shared a common ancestor dated circa 1961 (95% CI [1501.59, 1993.00]), and three of five isolates possessed a unique MDR profile (two isolates possessed no AMR genes; Figure 7).

The four other isolates sequenced in this study that were members of the ZA ST19/34 Clade were ST19 isolates, varied in their AMR profiles, and included: (i) POL_1967_GP_5765SD-B-S49 (isolated from South Africa's Gauteng province) and (ii) POL_1967_XX_7229SG-A-S29, both isolated from poultry feces in 1967 with no AMR determinants; (iii) PIG_2007_GP_1034ST-S15, which was isolated in 2007 from a swine organ in South Africa's Gauteng province and possessed tetracycline resistance gene *tet(B)*; (iv) POL_1985_XX_NEW25-S119, a MDR strain isolated in 1985 from poultry feces, which harbored nine AMR genes (streptomycin resistance genes *aadA4*, a*ph(3'')-Ib*, and *aph(6)-Id*, kanamycin resistance gene *aph(3')-Ia*, beta lactamase *blaTEM-1*, chloramphenicol resistance gene *catA1*, trimethoprim-resistance gene *dfrA21*, sulfonamide resistance gene *sul2*, and tetracycline resistance gene *tet(B)*; Figure 7).

An additional ST19 subclade within the ZA ST19/34 Clade did not contain any isolates sequenced in this study but contained ten genomes of MDR strains isolated in South Africa in 2001 (Figure 7). These genomes shared a common ancestor dated circa 1986 (95% CI [1861.51, 2001.0]), and each contained between 14 and 16 AMR genes; all ten genomes possessed gentamicin resistance gene *aac(3)-IIe*, streptomycin resistance genes *aadA1*, *aph(3'')-Ib*, and *aph(6)-Id*, rifamycin resistance gene *arr-2*, beta lactamases $bla_{SCO-1}$ and $bla_{TEM-131}$, (chloram)phenicol resistance genes *cmlA5* and *floR*, trimethoprim resistance gene *dfrA23*, sulfonamide resistance genes *sul1* and *sul2*, and tetracycline resistance gene *tet(A)* (Figure 7). Additional AMR genes that were variably present within this clade included (i) beta lactamase $bla_{SHV-5}$ (8 of 10 genomes, 80%); (ii) beta lactamase $bla_{OXA-17}$ (6 of 10 genomes, 60%); (iii) gentamicin resistance gene *aac(6')-Ib'*, (iv) amikacin/gentamicin/kanamycin/tobramycin resistance gene *aac(6')-Ib-AGKT*, (v) beta lactamase $bla_{OXA-10}$ (each detected in 4 of 10 genomes, 40%), and (vi) $bla_{SHV-165}$ (1 of 10 genomes, 10%; Figure 7).

**A largely MDR *S.* Typhimurium clade contains human- and animal-associated isolates from South Africa.** Three additional *S.* Typhimurium isolates sequenced here were members of a 22-isolate clade, which shared a common ancestor dated circa 1924 (95% CI [1413.97, 1967.00]; denoted in Figure 7 as "ZA Mixed MDR Clade"). Of the 22 genomes in this clade, 16 (72.7%) possessed between 4 and 7 AMR genes, while 6 genomes (27.3%) possessed no AMR genes (Figure 7). Two strains sequenced here (BOV_2005_GP_0892ST-S75, isolated in 2005 from bovine feces in Gauteng province, and XXX_2017_XX_89257-S68, isolated in 2017) possessed streptomycin resistance genes *aph(3'')-Ib* and *aph(6)-Id*, sulfonamide resistance gene *sul2*, and tetracycline resistance gene *tet(A)* (Figure 7). The third strain sequenced here (BOV_2015_XX_NEW16-S110, isolated in 2015 from bovine meat) possessed eight AMR genes: streptomycin resistance genes *aph(3'')-Ib, aph(6)-Id*, and *aadA1*, kanamycin resistance gene *aph(3')-Ia*, two full copies of beta lactamase $bla_{OXA-2}$, sulfonamide resistance gene *sul2*, and tetracycline resistance gene *tet(C)* (Figure 7). Other members of this clade included 14 genomes from humans in South Africa, one from a human in Tunisia, one from a human in Madagascar, one from bovine feces in Ethiopia, and one from each of Egypt and Senegal (Figure 7).

**A largely antimicrobial-susceptible clade with sporadic MDR contains South African and Nigerian subclades.** Three *S.* Typhimurium isolates sequenced in this study were contained within a 46-isolate clade, which shared a common ancestor dated circa 1913 (95% CI [1257.12, 1956.00]; denoted in Figure 7 as the "Nigerian/ZA Clade"). The three isolates clustered among seven publicly available South African genomes (six from humans and one environmental isolate) and shared a common ancestor dated circa 1965 (95% CI [1553.36, 1965.00]; Figure 7). While two of the publicly available genomes within this South African subclade possessed AMR gene profiles characteristic of MDR isolates, none of the eight remaining isolates possessed AMR genes, including the three isolates sequenced here; one strain (POL_1965_GP_7139SG-D-S62) was isolated from poultry feces in South Africa's Gauteng province in 1965, another (POL_2004_EC_9531ST-S78) from poultry meat in Eastern Cape in 2004, and a third (CAT_2003_KZ_9096ST-S8) from cat feces in KwaZulu Natal in 2003 (Figure 7). When compared to genomes from a previous study of *S.* Typhimurium in the United States (New York State, NYS) (Carroll et al., 2020), the three strains sequenced here formed their own clade, which did not contain any NYS genomes, and shared a common ancestor with a DT104-like clade (discussed in detail in the main text) circa 1816 (95% CI [1023.73, 1965.00]; Supplemental Figure S6).

**Three antimicrobial-susceptible *S.* Typhimurium isolates from animals in South Africa's Gauteng province shared a common ancestor with *S.* Typhimurium strains from camels in Egypt nearly 200 years ago.** Three additional *S.* Typhimurium isolates sequenced in this study clustered together, sharing a common ancestor dated circa 1956 (95% CI [1367.41, 2003.00]; denoted as part of the "Egyptian/ZA Clade" in Figure 7). All three were from South Africa's Gauteng province, and none possessed AMR genes (Figure 7): (i) PIG_2003_GP_12363-2ST-S76, isolated in 2003 from a swine organ, (ii) BOV_2006_GP_12565-4ST-S87, isolated in 2006 from bovine meat, and (iii) AVI_2008_GP_1844ST-S44, isolated in 2008 from pigeon feces. Members of this three-isolate clade shared a common ancestor with a seven-isolate clade dated circa 1836 (95% CI [1011.08, 1977.00]); the seven-isolate clade contained six Egyptian isolates (four isolated from camels in 2017, one isolated from basil in 2006, and one of unknown origin, isolated in 1977) and one from Madagascar (isolated from a human in 1979; Figure 7). Only one genome within the Egyptian/ZA Clade possessed AMR genes (i.e., the Egyptian strain isolated from basil in 2006, which possessed streptomycin resistance genes *aph(3")-Ib* and *aph(6)-Id*, sulfonamide resistance gene *sul2*, and tetracycline resistance gene *tet(A)*; Figure 7). The three Gauteng isolates sequenced here were not closely related to any of the *S.* Typhimurium strains from New York State (Supplemental Figure S6).

**A largely antimicrobial-susceptible clade primarily contains *S.* Typhimurium strains isolated from animals, food, and humans in Ethiopia.** Two additional *S.* Typhimurium isolates sequenced here were members of a 26-isolate clade, which shared a common ancestor dated to circa 1915 (95% CI [958.71, 1957.0]; denoted in Figure 7 as the "Ethiopian Clade"). No

AMR genes were detected in either OVI_1968_GP_8833-S38 (isolated in 1968 from ovine feces in South Africa's Gauteng province) and PIG_2004_GP_9239ST-S39 (also from Gauteng, isolated in 2004 from swine meat). Most isolates in this clade were from animals (i.e., from bovine, poultry, swine, and wild rodent sources), food (i.e., ground, cracked black pepper), and humans in Ethiopia (*n* = 18), with additional genomes from swine/unknown sources in Rwanda and poultry in Uganda (*n* = 2 each), and humans in South Africa (KwaZulu Natal) and the DRC (*n* = 1 each; Figure 7). The only genomes that possessed any AMR genes were the two poultry strains isolated in Uganda in 2016, which both possessed streptomycin resistance gene *aadA1*, beta lactamase *blaTEM-1*, and sulfonamide resistance gene *sul3* (Figure 7). The two South African isolates sequenced here were not closely related to any of the *S.* Typhimurium strains from New York State (Supplemental Figure S6).

**A largely antimicrobial-susceptible *S.* Typhimurium clade with sporadic AMR spans 13 African countries.** Another *S.* Typhimurium strain sequenced in this study (BOV_2001_XX_NEW20-S114, isolated in 2001 from bovine feces) was a member of a 39-isolate clade, which shared a common ancestor dated circa 1828 (95% CI [1082.50, 1956.00]; denoted in Figure 7 as the "Small Mixed Clade"). This clade encompassed strains from a variety of human-, animal-, environmental, and food-associated sources from Madagascar (*n* = 9), Morocco and Senegal (*n* = 7 each), South Africa (*n* = 5), Burkina Faso and Djibouti (*n* = 2 each), and Guinea, Algeria, Cameroon, Ethiopia, Malawi, Chad, and Nigeria (*n* = 1 each). Of all 39 genomes within the Small Mixed Clade, 32 (82.1%) possessed no AMR genes. The remaining genomes possessed between one and nine AMR genes; the strain sequenced here possessed one AMR gene, tetracycline resistance gene *tet(A)* (Figure 7). When compared to bovine- and human-associated strains from New York State, the bovine isolate sequenced here clustered among 40 New York State isolates from humans, dairy cattle, and dairy farm environments (Supplemental Figure S6), which were largely antimicrobial susceptible, but showcased sporadic AMR acquisition (Carroll et al., 2020). All 41 isolates shared a common ancestor dated circa 1848 (95% CI [1187.65, 2001]; Supplemental Figure S6).

**An antimicrobial-susceptible South African strain shares a most recent common ancestor with a MDR Senegalese strain.** Another *S.* Typhimurium strain sequenced in this study (BOV_1990_XX_NEW18-S112, isolated in 1990 from bovine feces; denoted in Figure 7 with a purple arrow) possessed no AMR genes. This South African strain was most closely related to a strain isolated from an unknown source in Senegal in 1999; the Senegalese strain possessed four AMR genes: streptomycin resistance genes *aph(3'')-Ib* and *aph(6)-Id,* sulfonamide resistance gene *sul2*, and tetracycline resistance gene *tet(A)* (Figure 7). These two strains shared a common ancestor dated circa 1955 (95% CI [1525.31, 1990.00]; Figure 7). The South African isolate sequenced here was not closely related to any of the *S.* Typhimurium strains from New York State (Supplemental Figure S6).

**A largely antimicrobial-susceptible South African clade primarily contains human-associated isolates.** The remaining *S.* Typhimurium strain sequenced in this study (POL_1967_GP_2671SG-S83, isolated in 1967 from poultry feces in South Africa's Gauteng province) possessed no AMR genes and was a member of an 18-isolate clade with a common ancestor dated circa 1920 (95% CI [1409.02, 1959.00]; denoted in Figure 7 as the "ZA Small Clade"). This clade primarily contained human isolates from South Africa ($n$ = 14), as well as an isolate from each of Madagascar (human isolate), Mali (human isolate), and Burkina Faso (unknown isolate; Figure 7). Within the ZA Small Clade, only one strain, isolated in 2020 from a human in South Africa's Gauteng province, possessed AMR genes: beta lactamase *blaTEM-1* and fosfomycin resistance gene *fosA3* (Figure 7). The South African isolate sequenced here was not closely related to any of the *S.* Typhimurium strains from New York State (Supplemental Figure S6).

**Supplemental Text References**


Carroll, L.M., Huisman, J.S., and Wiedmann, M. (2020). Twentieth-century emergence of antimicrobial resistant human- and bovine-associated *Salmonella enterica* serotype Typhimurium lineages in New York State. *Sci Rep* 10**,** 14428.
